# NF-κB-Inducing Kinase Maintains Mitochondrial Efficiency and Systemic Metabolic Homeostasis

**DOI:** 10.1101/2021.08.26.457753

**Authors:** Kathryn M. Pflug, Dong W. Lee, Justin Keeney, Raquel Sitcheran

## Abstract

**Background:** NF-κB-inducing kinase (NIK) is a critical regulator of immunity and inflammation and NIK loss-of-function mutations have recently been described in patients with primary immunodeficiency disease. Based on our previous work showing that NIK regulates adaptive metabolic responses in glucose-starved cancer cells, we investigated whether NIK is required for mitochondrial functions in bioenergetic processes and metabolic responses to nutritional stress in NIK knockout (KO) mice, which recapitulate the clinical presentation of NIK PID patients.

**Methods:** We performed whole body composition analysis of wild type (WT) and NIK KO mice using EchoMRI and DEXA imaging. Seahorse extracellular flux analyses were used to monitor oxidative phosphorylation and glycolysis through oxygen consumption rates (OCR) and extracellular acidification rates (ECAR) in preadipocyte cells and in ex vivo adipose tissue. NIK regulation of systemic metabolic output was measured by indirect calorimetry using TSE Phenomaster metabolic chambers under basal conditions as well as in response to nutritional stress induced by a prolonged high-fat diet (HFD). Finally, we analyzed a role for NIK in adipocyte differentiation, as well as the contributions of canonical and noncanonical NF-κB signaling to adipose development and metabolic output.

**Results:** We observed that in adipose cells, NIK is required for maintaining efficient mitochondrial membrane potential and spare respiratory capacity (SRC), indicators of mitochondrial fitness. NIK KO preadipocytes and ex vivo adipose tissue exhibited diminished SRC, increased proton leak, with compensatory upregulation of glycolysis. Systemically, NIK KO mice exhibited increased glucose utilization, increased energy expenditure, and reduced adiposity, which persisted under the stress of HFD. Finally, while NIK controlled adipocyte differentiation through activation of RelB and the noncanonical NF-κB pathway, NIK regulation of metabolism in preadipocytes was NF-κB/RelB-independent.

**Conclusion:** Our results demonstrate that NIK is required for metabolic homeostasis both locally, on a cellular and tissue level, as well as systemically, on an organismal level. Collectively, the data suggest that NIK KO cells upregulate glycolytic metabolism as a compensatory response to impaired mitochondrial fitness (diminished SRC) and mitochondrial efficiency (increased proton leak). To meet changes in bioenergetic demands, NIK KO mice undergo metabolic rewiring through increased glucose utilization and glycolysis, which persists under the stress of overnutrition with a HFD. Moreover, while NIK regulation of metabolism is RelB-independent, NIK regulation of adipocyte development requires RelB and activation of the noncanonical NF-κB pathway. Our findings establish NIK as an important regulator of cellular and systemic metabolic homeostasis, suggesting that metabolic dysfunction may be an important component of primary immunodeficiency diseases arising from loss of NIK function.

## 1. Introduction

Nuclear Factor-κB (NF-κB)-inducing kinase (NIK), encoded by *MAP3K14*, is a serine, threonine protein kinase that that is best known for its function as an upstream inducer of NF-κB signaling to regulate innate and adaptive immunity. While NIK can activate canonical and noncanonical NF-κB pathways, it is uniquely required for activation of noncanonical NF-κB signaling through phosphorylation of Inhibitor of κB Kinase α (IKKα), which triggers p100 processing to p52, and generation of transcriptionally active p52-RelB NF-κB complexes [1–4]. NIK has critical roles in B-cell, lymphocyte, and lymph node development, immunoglobulin (Ig) production and T-cell function. Consequently, *Nik^aly^* mutant mice (*aly*; alymphoplasia), which lack NIK activity, and NIK knockout mice (NIK KO) exhibit lymphopenia, abnormal Peyer’s patches, aberrant splenic and thymic structures, reduced B-cell numbers and Ig serum levels leading to humoral immunodeficiency [1,5–9]. In humans, NIK loss-of-function mutations were recently identified in patients with primary immunodeficiency (PID) who exhibit similar immune defects as *Nik^aly^* and NIK KO mice [10, 11], demonstrating the relevance of NIK deficient mouse models for immunodeficiency disease.

We, and others, have elucidated important NF-κB-independent metabolic functions for NIK, including roles in mitochondrial dynamics, metabolic reprogramming in macrophages and under nutrient stress in cancer cells, as well as regulation of glycolysis in T-cells [12–15]. In response to diet-induced obesity, gain-of-function studies have shown that NIK induces hyperglycemia by increasing glucagon activity, and liver steatosis through induction of fatty acid oxidation, and loss of NIK was shown to increase glucose and insulin tolerance [16–18]. Our recent work has established NIK mitochondrial localization and regulation of mitochondrial respiration and fitness, promoting metabolic adaptation of cancer cells to glucose starvation [15]. However, NIK regulation of mitochondrial metabolism has not been examined in adipose tissue, or on an organismal level.

Metabolic homeostasis relies on a balance between inputs such as glucose and oxygen necessary for energetic processes and utilization of catabolic pathways involving glycolysis and oxidative phosphorylation (OXPHOS) for efficient energy production [19, 20]. Utilization of these major metabolic pathways is dynamic and shifts in response to alterations in cellular states or stress [21, 22]. Glycolysis and OXPHOS are interwoven and dysregulation in one can alter the flux or rate of the other [23–25]. Although glycolysis yields low amounts of ATP, it is important for feeding into the TCA cycle and electron transport chain (ETC) for rapid production of ATP, particularly under low oxygen availability [26–28]. Furthermore, glucose catabolism is preferred in response to stress as it requires the lowest input of ATP [29–31]. Oxidative phosphorylation, or aerobic respiration, is more commonly utilized to meet ATP demands. Important for mitochondrial respiration is its spare respiratory capacity (SRC), a measurable indicator of mitochondrial fitness, or the ability to upregulate oxygen consumption and ATP production to meet changes in energetic demand [32–34]. Energy derived from mitochondria is essential for maintaining cellular homeostasis, as well as in response to stress. Reciprocally, on an organismal level, alterations in mitochondrial functions impact physiological and behavioral responses, as seen in individuals with impaired ETC function due to primary mitochondrial disorders (PMDs), who are unable to adapt to dietary and physiological stressors [35, 36]. As such, mitochondrial fitness and efficiency is essential to organismal homeostasis and adaptation to bioenergetic stress.

Here we investigate a functional and developmental role for NIK in controlling metabolic homeostasis basally and in response to nutritional stress induced by a high-fat diet (HFD). Our data demonstrate that NIK deficient preadipocytes and *ex vivo* tissue exhibited loss of mitochondrial SRC and increased proton leak. This uncoupling of mitochondrial respiration and inefficient metabolism was accompanied by a compensatory increase in glycolysis and extracellular acidification rate (ECAR). Furthermore, we show that NIK regulation of mitochondrial SRC and proton leak is independent of noncanonical RelB/NF-κB signaling. Similar to NIK KO cells and tissue, NIK deficiency in mice resulted in increased glucose utilization and glycolysis to meet energetic demands. In NIK KO mice, and *ex vivo* adipose tissue, this increased energy expenditure resulted in reduced adiposity and resistance to adipose accumulation and weight gain through development, as well as under a HFD. While NIK regulates mitochondrial functions independently of RelB/NF-κB, we found that NIK promotes adipocyte differentiation in a noncanonical NF-κB-dependent manner through transcriptional regulation of key adipogenic transcription factors. These results are the first to describe NIK regulation of adiposity through NF-κB-dependent development signaling, as well as NF-κB-independent regulation of local-tissue metabolism and systemic energy expenditure, suggesting that systemic mitochondrial metabolic defects are an underlying component of immune dysfunction.

## 2. Results

### 2.1 NIK KO mice have reduced adiposity

Whole body composition analyses were performed using dual-energy x-ray absorptiometry (DEXA) imaging to evaluate potential alterations in systemic body composition due to NIK deficiency. Mice were utilized between the ages of 2-4 months, that exhibited a healthy deposition. We observed that NIK KO mice displayed a significant reduction in overall fat mass and considerably smaller visceral and subcutaneous adipose tissue depots (Figure 1A). Further assessment of whole body composition using EchoMRI^TM^, demonstrated that both NIK KO males and females had a significant reduction in overall fat mass and a higher lean mass ratio relative to body weight (Figure 1B,C). Visceral gonadal white adipose tissue (gWAT) depots were substantially reduced in male and female NIK KO mice compared to WT and HET mice (Figure 1D,E, Supp. Figure 1A). Similarly, subcutaneous inguinal white adipose tissue (ingWAT) was reduced in NIK KO mice (Supp. Figure 1B). While WT mice increased in fat mass as they aged from 2 to 4 months, NIK KO mice maintained a reduced fat mass during this time period (Figure 1F, Supp. Figure 1C,D), and exhibited no signs of infection. Although NIK KO mice gained less fat, this was not due to behavioral changes as they consumed more food and water with reduced physical activity than their WT littermates (Supp. Figure 1C,D). Furthermore, though a reduction in adiposity was notable even at 2 months age (Supp. Fig. 1F), NIK KO mice did not significantly differ in weight compared to WT mice until 15 weeks of age in males, possibly when the rate of fat mass development in WT mice surpassed the rate of body growth and adipose development in NIK KO mice (Supp. Figure 1G,H). Oil Red O staining of liver sections revealed no differences in exogenous liver lipid accumulation between WT and NIK KO mice (Supp. Figure 1I), which can occur due to defects in adipocyte function and lipolysis [37, 38], suggesting a role for NIK in the adipose tissue development.

**Figure 1:**
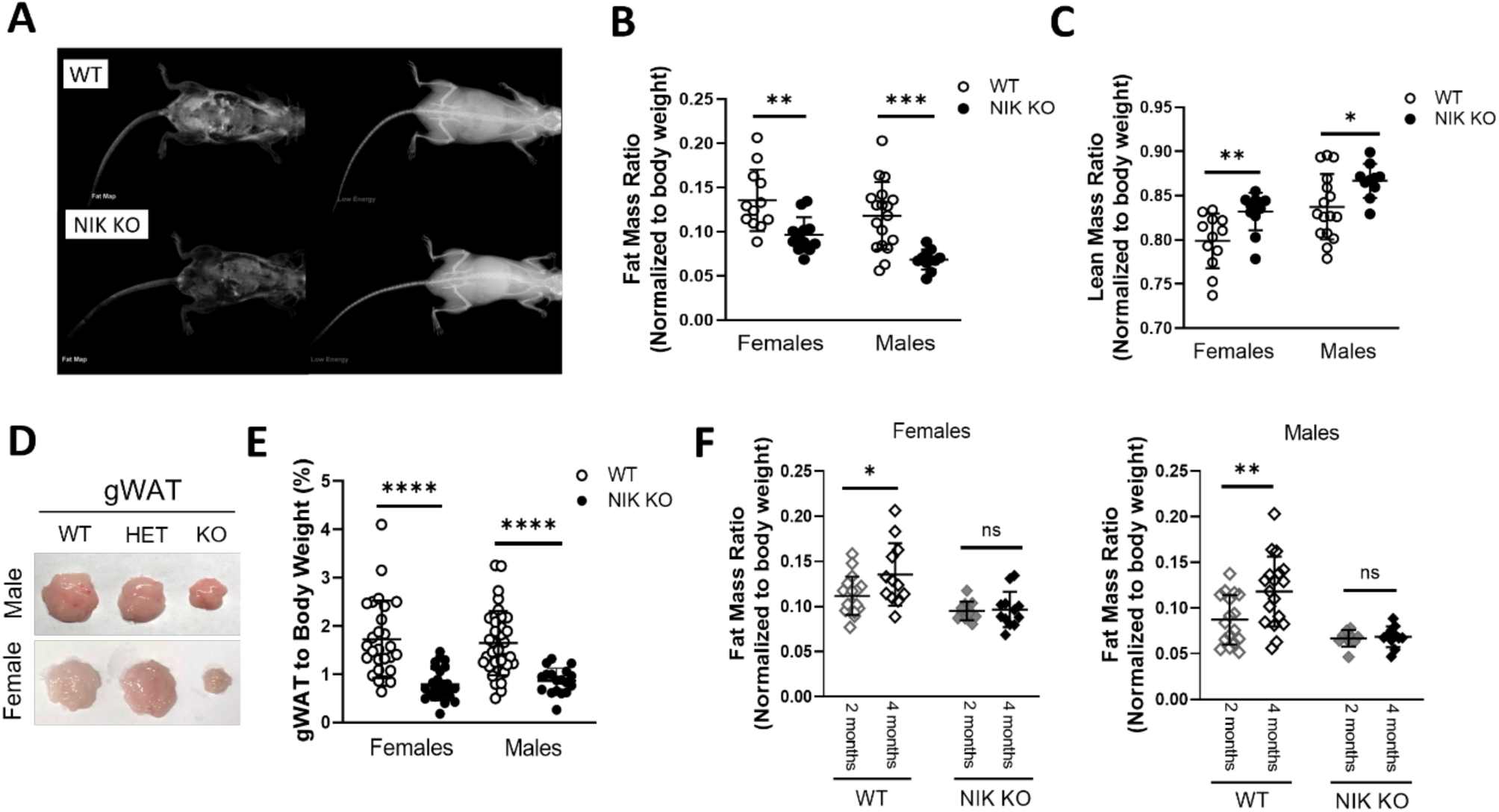
NIK KO mice have reduced adiposity. **(A)** DEXA scans of chow fed male WT and NIK KO mice at 2 months, displaying overall fat map (left) and overall low energy map (right). **(B)** EchoMRI^TM^ measurements of overall fat and **(C)** lean mass of male and female mice normalized to body weight. WT n= 12 females and 18 males, KO n=12 females and 11 males. **(D)** Gonadal adipose tissue from male or female mice. **(E)** Weight of gonadal fat between WT and NIK KO male and female mice normalized to body weight. WT n= 27 females and 36 males, KO n= 26 females and 18 males. **(F)** Fat mass ratio from body composition data of female and male WT and NIK KO mice at 2 and 4 months of age. At 2-months-old WT n=17 females and 17 males, KO n= 9 females and 9 males. At 4-months-old WT n= 12 females and 18 males, KO n= 12 females and 11 males. **(B-C,E-F)** Data represented as mean ± SD, Unpaired Student t-test.

### 2.2 NIK regulates mitochondrial respiration and efficiency

We previously demonstrated that NIK localizes to mitochondria, promoting mitochondrial fission and oxidative metabolism in glioblastoma cells [13]. Similarly, we observed that multipotent C3H10T1/2 cells, capable of differentiating into adipocytes, also exhibited larger, more fused mitochondria when NIK expression was ablated. Exogenous expression of human NIK in NIK KO cells (NIK KO-hNIK) restored mitochondrial morphology similar to that observed in WT cells (Figure 2A,B). Similarly, 3T3-L1 NIK KO preadipocytes also exhibited a fused mitochondria phenotype. However, cells lacking the downstream noncanonical NF-κB transcription factor, RelB (RelB KO cells), did not exhibit a fused mitochondrial morphology similar to NIK KO cells, suggesting this phenotype in is NIK-dependent (Figure 2C). Given the reduction in adiposity and changes in mitochondria morphology in NIK KO cells, we sought to determine whether NIK regulates mitochondrial function in adipocytes. Metabolic analysis of 3T3-L1 cells by Seahorse extracellular flux assay revealed that NIK KO cells exhibited higher basal oxygen consumption rate (OCR) but with significantly impaired mitochondrial spare respiratory capacity (SRC), and increased mitochondrial proton leak, characteristic of inefficient mitochondrial metabolism (Figure 2D). Normal SRC and proton leak phenotypes were restored in NIK KO-hNIK cells (Figure 2D, Supp. Fig 1J), demonstrating that the aberrant mitochondrial metabolism is NIK-dependent. RelB KO cells did not have impaired SRC and proton leak, showing that NIK regulation of mitochondrial efficiency is also noncanonical NF-κB-independent (Figure 2D). NIK KO and NIK KO-hNIK cells also showed an increase in basal extracellular acidification rate (ECAR), a readout for glycolysis, while RelB KO cell ECAR was similar to WT (Figure 2E), consistent with NF-κB-independent roles for NIK in adipocyte metabolism. Basally, NIK KO, NIK KO-hNIK and RelB KO cells had similar basal energetic profiles compared with WT cells. However, NIK KO cells exhibited impaired OCR in response to mitochondrial stressors, which was accompanied by a compensatory increase in glycolysis, whereas RelB KO cell metabolism was unaffected (Figure 2F). Exogenous NIK expression rescued OCR elevation under a stressed state in NIK KO cells, demonstrating a requirement for NIK to meet metabolic demand in a stressed state through OXPHOS (Figure 2G).

**Figure 2:**
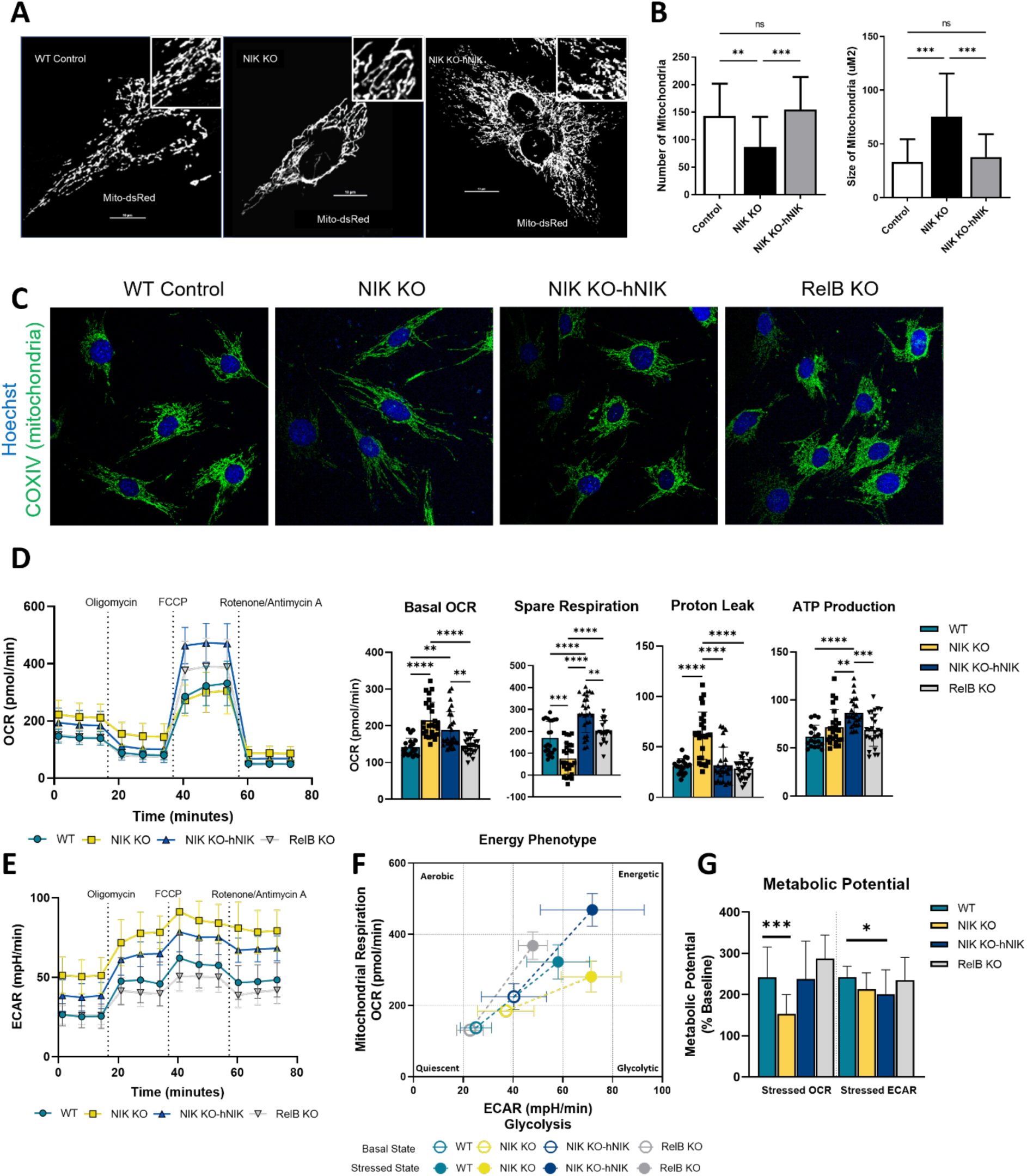
NIK regulates mitochondrial respiration and efficiency. **(A)** Representative imaging of C3H10T1/2, mouse stem cells, transfected with Mito-dsRed. **(B)** Image J quantification of Mito-dsRed transfected WT, NIK KO, and NIK KO-hNIK C3H10T1/2 cells. Data represented as mean ± SD, One-Way ANOVA **(C)** Immunofluorescence imaging of WT, NIK KO, NIK KO-hNIK, and RelB KO C3H10T1/2 cells with mitochondria (COXIV, green) and nuclear (Hoechst, blue) staining. **(D)** Seahorse extracellular flux analysis of 3T3-L1 preadipocytes oxidative stress test (mitochondria stress test) displaying oxygen consumption rate (OCR) (statistics compared to WT) and **(E)** extracellular acidification rate (ECAR). **(D-E)** OCR and ECAR line graphs are represented as mean ± SEM. Mitochondria stress test dot plots are represented as mean ± SD, One-Way ANOVA. N= 4 independent, biological replicates with 5-8 technical replicates per independent run. **(F)** Cell energy phenotype of 3T3-L1 cells from seahorse analysis showing basal states of cell (open square) by OCR and ECAR to stressed states (closed square). Data represented as mean ± SD. **(G)** Metabolic potential of 3T3-L1 cells from seahorse analysis with contribution by either OXPHOS (OCR) or by glycolysis (ECAR) after FCCP injection (stressed rates). Data represented as mean ± SD, One-Way ANOVA**. (F-G**). N= 3 independent, biological replicates with 5-8 technical replicates per independent run.

### 2.3 Glycolytic dependence increases in response to metabolic demand in NIK KO adipose tissue

To evaluate whether NIK regulates adipocyte metabolism in the context of its native microenvironment, we analyzed OCR and ECAR in *ex vivo* adipose tissue from WT and NIK KO mice. Although basal OCR was not significantly altered, NIK KO ingWAT and gWAT depots exhibited impaired maximal respiration and reduced SRC, increased proton leak and decreased ATP production, similar to adipocyte cells (Figure 3A). We also observed robustly elevated ECAR in both NIK KO adipose depots, consistent with metabolic reprogramming to meet energetic demands and compensate for inefficient OXPHOS (Figure 3B). Both adipose depots were able to increase glycerol levels in response to isoproterenol, a β-agonist and lipolytic stimulant, suggesting that NIK KO adipose tissues are functional, with gWAT displaying enhanced lipid turnover compared to WT (Figure 3C). Together, these findings suggest that loss of NIK in adipocytes and adipose tissue resulted in mitochondrial dysfunction exhibited by impaired SRC and elevated proton leak, causing an increase in energetic demand that is met by enhanced glycolytic metabolism.

**Figure 3:**
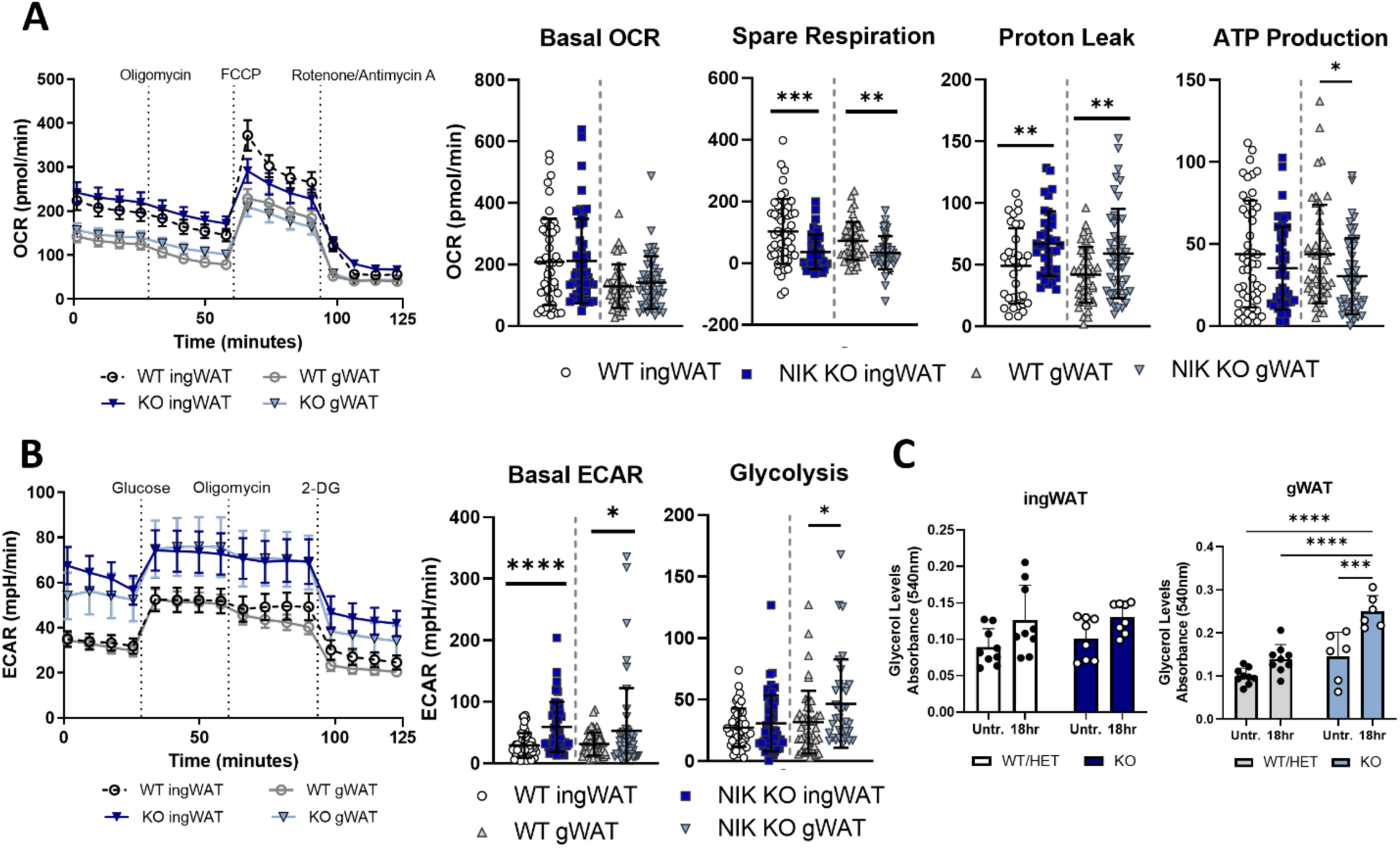
Glycolytic dependence increases in response to metabolic demand in NIK KO mice. **(A)** Seahorse extracellular flux analysis of *ex vivo* mouse gonadal (gWAT) or inguinal WAT (ingWAT) with an oxidative stress test (Mitochondria Stress Test). **(B)** Seahorse extracellular flux analysis of *ex vivo* mouse gonadal or inguinal WAT with Glycolysis Stress Test. **(A-B)** Data represented as mean ± SEM for line graphs and mean ± SD for dot plots. WT n=4 females and 5 males, KO n= 4 females and 3 males. Data normalized by protein. **(C)** Glycerol levels of gonadal or inguinal WAT with lipolysis stimulated with isoproterenol and measured prior to treatment (untreated) and after 18-hour incubation. Data represented as mean ± SD, Sidak’s Multiple Comparison Test. Data normalized by tissue weight.

### 2.4 Sex dimorphic metabolic differences are exhibited in NIK KO mice

We next sought to evaluate whether NIK-dependent metabolism observed in cellular and adipose tissue impacted systemic metabolism. Using indirect calorimetry metabolic cages, whole body analyses of WT and NIK KO mice were performed across light and dark cycles to analyze respiratory rates and energy expenditure, along with feed and water intake as well as activity. Overall, female NIK KO mice displayed higher oxygen consumption and carbon dioxide production rate throughout both periods of the day compared to their WT counterparts, while NIK KO male systemic metabolic rates were not significantly altered (Figure 4 A, B). By measuring respiration exchange rates (RER), substrate utilization for oxidation can be estimated. Glucose oxidation has the highest respiration exchange rate, followed by protein, and then lipid oxidation [29]. Tracking changes in oxidation preference between inactive (light) and active (dark) cycles by RER measurements revealed that at 2 months of age, both WT and NIK KO mice similarly utilize lipid oxidation during their inactive states (Supp. Figure 2D). However, as NIK KO mice age, glucose or protein utilization for oxidation increases during their inactive periods, similar to RER in their active periods. In contrast, WT mice maintain lower RER in their inactive states, indicative of lipid oxidation. During active periods, WT and NIK KO RER increase similarly at both ages (Figure 4C). Moreover, NIK KO mice displayed a higher overall energy expenditure, particularly in females compared with males (Figure 4D). In addition to increased metabolic output, NIK KO mice also exhibited higher temperatures at 2 months of age (Figure 4E), and female mice maintained elevated temperatures at 4 months (Supp. Figure 2A). Consistent with increased temperature and energy expenditure, we observed higher brown adipose tissue in both female and male NIK KO mice (Figure 4F). While male and female mice displayed adipose specific increases in metabolism from *ex vivo* extracellular flux analysis, only female mice exhibited systemic differences on a chow diet. This trend for an overall increase in whole body metabolic activity in the NIK KO mice is observable in aging mice (Supp. Figure 2).

**Figure 4:**
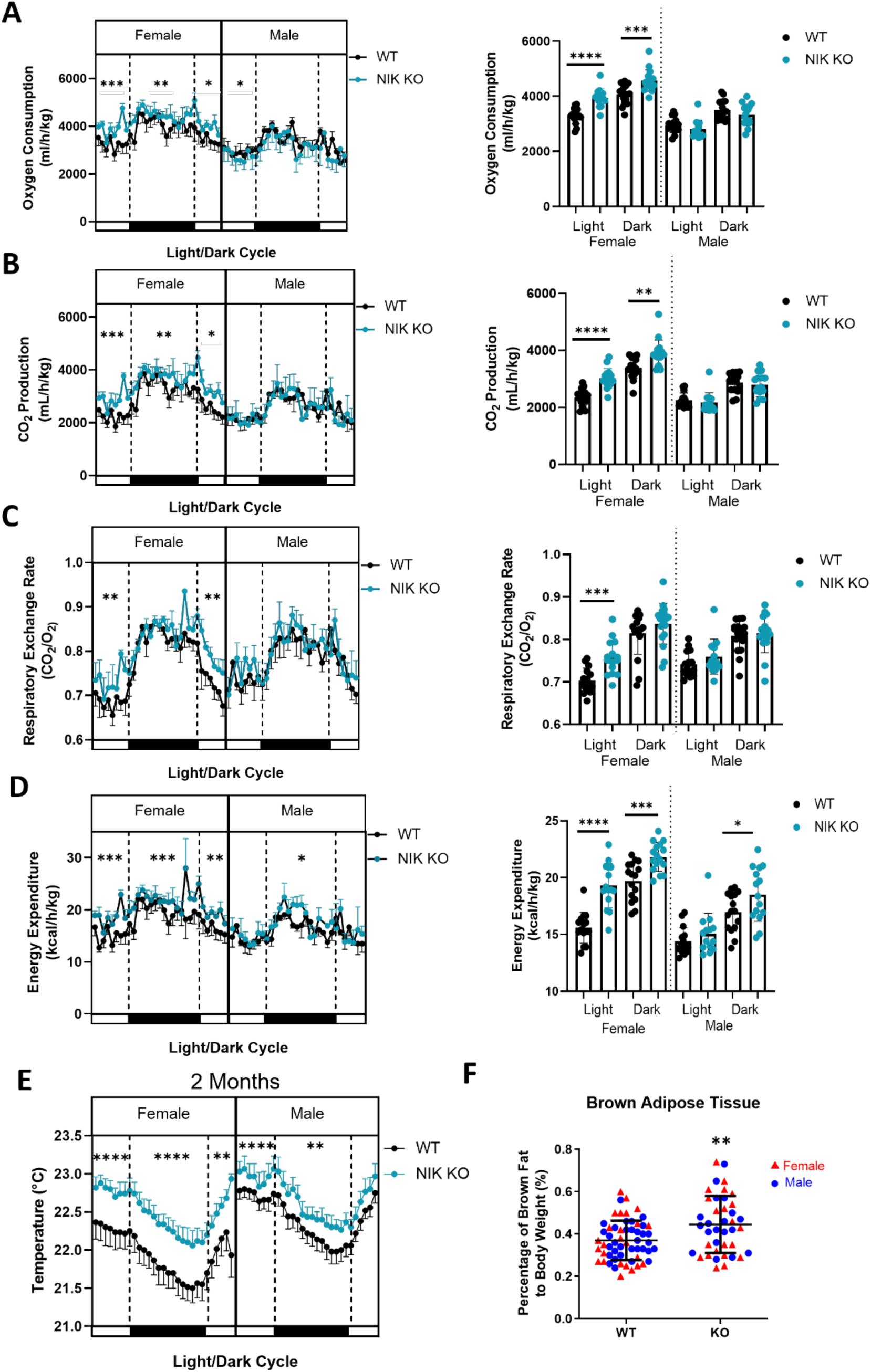
Sex dimorphic metabolic differences are exhibited in NIK KO mice. **(A-D)** Systemic metabolic data collected from 4-month-old, chow fed male and female *Map3k14* mice individually housed in TSE Phenomaster metabolic cages. Morning (light) and night (dark) analysis from a 24-hour time period of **(A)** oxygen consumption, **(B)** CO_2_ production, **(C)** respiratory exchange rate (CO_2_/O_2_), and **(D)** caloric energy expenditure. **(E)** Metabolic cage temperature with mice at 2 months of age. **(F)** Brown adipose tissue weights from male and female, chow fed WT and NIK KO mice at 4 months of age. Data represented as mean ± SD Males; WT n=25 females and 33 males, KO n=22 females and 19 males. Line graphs represented as mean ± SEM, bar graphs represented as mean ± SD. **(A-D)** Females WT n=5 females and 4 males, KO n=6 females and 3 males. **(E)** WT n= 6 females and 5 males, KO n=5 females and 3 males. **(A-F)** Data analyzed by Unpaired Student t-test.

### 2.5 NIK promotes adiposity in response to overnutrition with a high-fat diet

Next, we investigated whether NIK plays a role in regulating systemic metabolism in response to nutritional stress. After 2-3 months on a high-fat diet (HFD), both male and female NIK KO mice exhibited significantly less weight gain than their WT counterparts (Figure 5A, B, C). Whole body composition analysis by EchoMRI demonstrated that NIK KO mice maintained lower fat mass and a higher lean mass ratio by body weight compared to WT mice (Figure 5D, E). Overall, NIK KO mice gained about 3x less fat than WT mice on a HFD compared with standard chow (Figure 5F). Analysis of individual adipose depots revealed that NIK KO mice on a HFD exhibited a 50% decrease in gWAT tissue, along with a significant reduction in the subcutaneous depot, ingWAT, compared to WT mice (Figure 5G,H). In addition to exhibiting reduced adiposity, NIK KO mice were more efficient at clearing glucose from the blood in glucose tolerance tests compared to WT littermates, suggesting an increase in insulin sensitivity (Figure 5I). Insulin tolerance testing confirmed this, demonstrating that NIK KO mice also cleared glucose more rapidly than WT mice after an administration of insulin (Figure 5J). Analysis of adipogenic gene expression from the inguinal WAT of HFD mice also revealed a reduction in fatty acid binding protein 4 (*FABP4*; Ap2), and an increase in GLUT4 and *GPD1* (Supp. Figure 3A,B), which is consistent with both an increase in glucose clearance of NIK KO mice and higher glycolytic demands of NIK KO adipose tissue. Moreover, glycogen synthase expression in the livers of NIK KO mice is decreased, suggesting reduced glucose storage compared to WT (Supp. Figure 3C,D). NIK KO mice on a HFD also exhibited reduced lipid levels in the liver compared to WT mice accompanied by lower gene expression of fatty acid binding protein (*FABP1*) and fatty acid oxidation genes (*MCAD, CPT1α*) (Supp. Figure 3D,E). Overall, these data demonstrated the critical role NIK has in facilitating adipose expansion and glucose homeostasis with overnutrition under a HFD.

**Figure 5:**
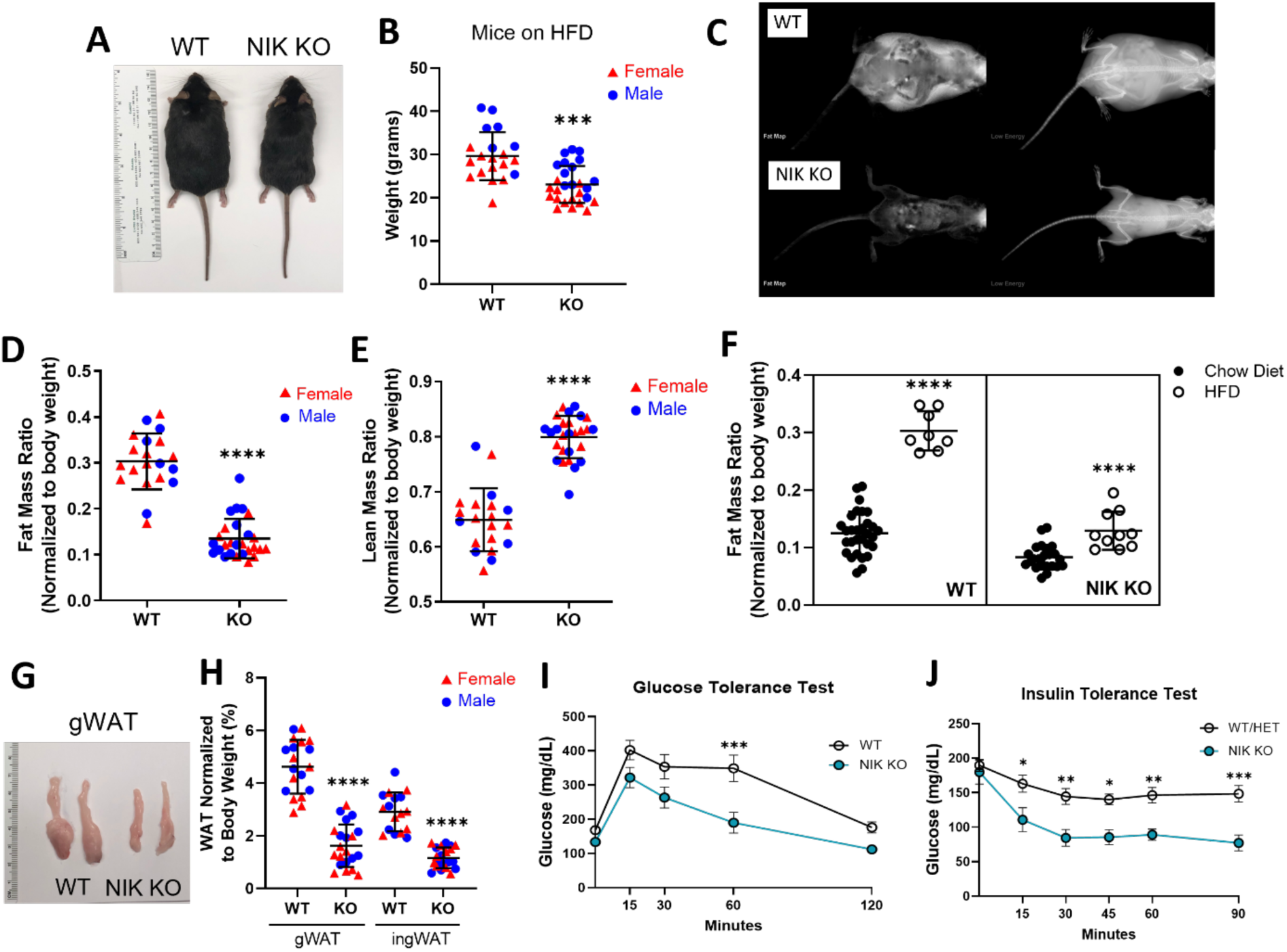
NIK promotes adiposity in response to overnutrition with a high-fat diet. **(A)** Images of male WT and NIK KO mice on HFD. **(B)** Weights of WT and NIK KO male and female mice exposed to a HFD for about 3 months (mice exposed to a HFD in utero up until weaning and then 8-10 weeks after weaning). Data represented as mean ± SEM, Unpaired Student t-test. **(C)** DEXA images of WT and NIK KO male mice on a HFD at about 3 months of age, displaying overall fat map (left) and overall low energy map (right). **(D)** EchoMRI^TM^ data of fat and **(E)** lean mass from female and male WT and NIK KO mice on a HFD normalized to weight. Data represented as mean ± SD, Unpaired Student t-test. **(F)** Fat mass data from EchoMRI^TM^ analysis comparing fat mass ratios from a chow diet to a HFD between WT and NIK KO mice. **(G)** Gonadal white adipose tissue from male mice on a HFD. **(H)** Weights of gonadal and inguinal white adipose tissue from male and female mice normalized to body weight. Data represented as mean ± SD, Unpaired Student t-test. **(I)** Glucose tolerance test on WT and NIK KO mice on a HFD. Data represented at mean ± SEM, Sidak’s Multiple Comparisons Test. WT n=8, NIK KO n=12. **(J)** Insulin tolerance test on WT, HET and NIK KO mice on a HFD. Data represented at mean ± SEM, Sidak’s Multiple Comparisons Test. WT/HET n=10, NIK KO n=6.

### 2.6 NIK is required for metabolic homeostasis in response to chronic dietary stress

Extracellular flux analyses were then performed on *ex vivo* adipose tissue from mice subjected to the chronic overnutrition with a HFD in utero through the age of 3 months. Adipose tissue from NIK KO mice on a HFD showed a striking increase in basal OCR and ATP production (Figure 6A). NIK KO ingWAT had significantly decreased SRC, while proton leak was consistently increased in both NIK KO depots (Figure 6A), suggesting elevated, but inefficient, mitochondrial metabolism, similar to results observed in NIK KO preadipocytes and *ex vivo* NIK KO adipose tissue from chow-fed mice. Analysis of glycolytic metabolism also demonstrated similar results to chow-fed mice, with NIK KO adipose tissue exhibiting significantly higher basal ECAR rates and higher levels of glycolysis (Figure 6B). We also observed elevated systemic metabolism in NIK KO mice on a HFD. Although we observed some sex dimorphic metabolic dysregulation in NIK KO mice under a chow diet, with the chronic stress of a HFD, both male and female NIK KO mice displayed elevated oxygen consumption and CO_2_ production (Supp. Fig. 4A,B) along with higher respiratory exchange rates, particularly during the light cycle, or inactive period (Figure 6C, Supp. Fig 4C). Notably, robust increases in energy expenditure and cage temperature were observed in NIK KO mice, demonstrating considerably elevated metabolic output compared to WT mice at all times of the day with HFD overnutrition (Figure 6D, Supp. Fig. 4D,E). Notably, leaner NIK KO mice did not exhibit increased physical activity (Supp. Fig. 4F), or reduced water or feed intake (Supp. Fig. 4G,H). Overall, our findings revealed that under either basal or nutrient stressed conditions NIK KO mice exhibited elevated metabolism with higher energy expenditure but have inefficient mitochondrial metabolism through impaired SRC and increased proton leak. Furthermore, increased glycolysis and glucose utilization persisted to meet energetic demands under the stress of a HFD, demonstrating an important role for NIK in metabolic homeostasis.

**Figure 6:**
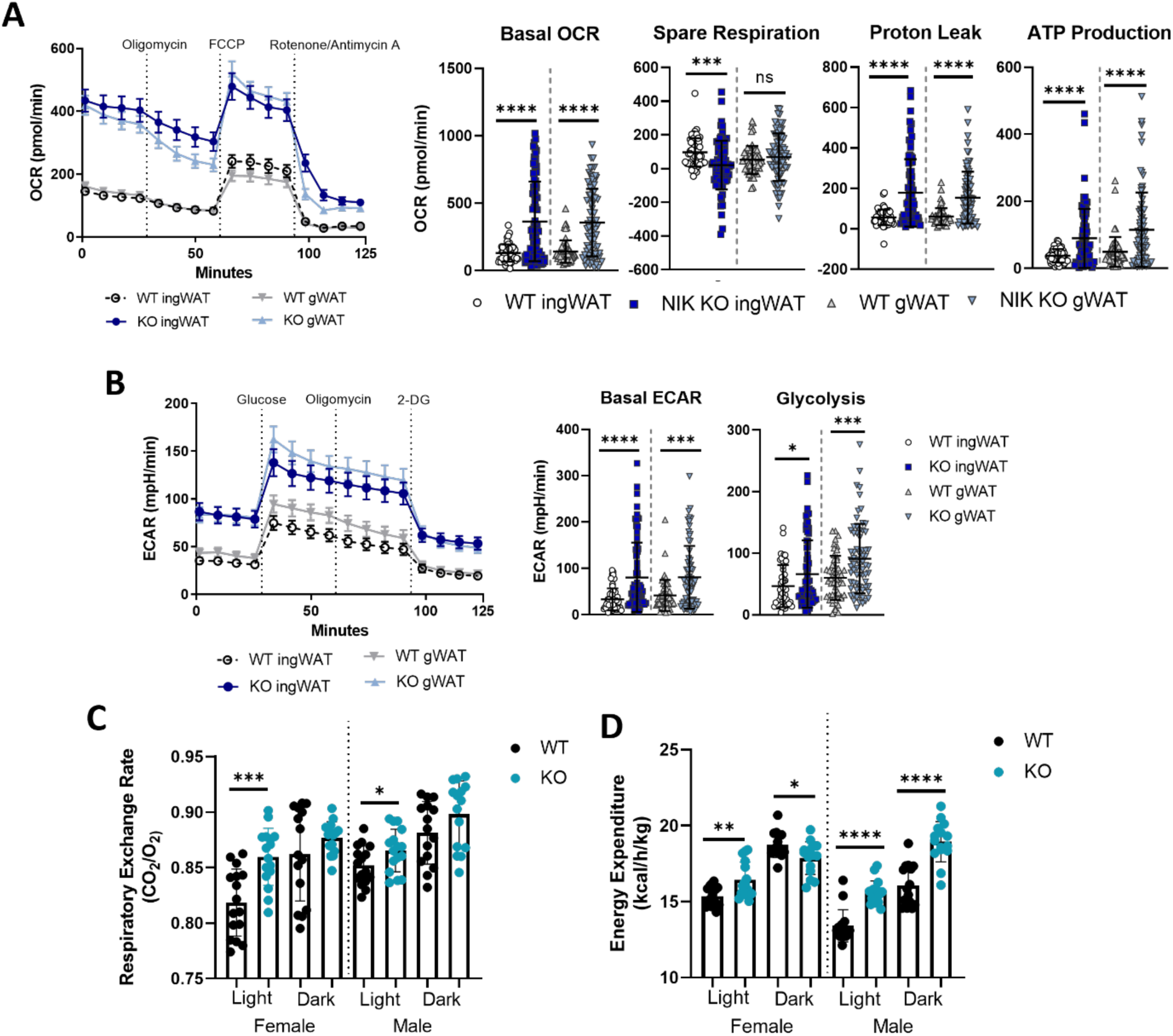
NIK is required for metabolic homeostasis in response to chronic dietary stress. **(A)** Seahorse extracellular flux analysis of *ex vivo* mouse gonadal or inguinal WAT with either oxidative stress test (Mitochondria Stress Test) or with **(B)** Glycolysis Stress Test. **(A,B)** Data represented as mean ± SEM for line graphs and mean ± SD for dot plots, Unpaired Student t-test. WT n=4 females and 5 males, KO n=6 females and 7 males. **(C,D)** Indirect calorimetry data collected from TSE Phenomaster cages of individually housed male and female *Map3k14* mice on a HFD at about 3 months of age. Morning and night analysis from a 24-hour time period of **(C)** respiratory exchange rate (CO_2_/ O_2_), and **(D)** caloric energy expenditure. WT n= 7 females and 5 males, KO n=8 females and 5 males. Data represented as mean ± SD, Unpaired Student t-test.

### 2.7 A role for noncanonical NF-κB-dependent signaling in promoting adiposity

Given previous studies suggesting a role for NF-κB in adipocyte differentiation and development, we investigated whether NIK-dependent noncanonical NF-κB activity was involved in promoting adiposity. Compared to control cells, NIK KO C3H10T1/2 and 3T3-L1 cells were severely impaired in their ability to form mature adipocytes, as demonstrated by staining of lipid droplets. Expression of a wild-type human NIK construct in NIK KO cells (NIK KO-hNIK) restored adipogenesis to the same extent as control cells (Figure 7A, Supp. Figure 5A,B). We also observed that primary bone marrow-derived stem cells (MSCs) isolated from NIK KO mice were impaired in their ability to differentiate into mature adipocytes compared to MSCs isolated from WT mice (Supp. Figure 5C). Furthermore, treatment of WT C3H10T1/2 cells with a NIK-specific inhibitor, B022, impeded adipocyte differentiation similar to cells lacking NIK (Supp. Figure 5D,E) [39]. These results support a crucial, cell-intrinsic role for NIK in adipocyte development.

**Figure 7:**
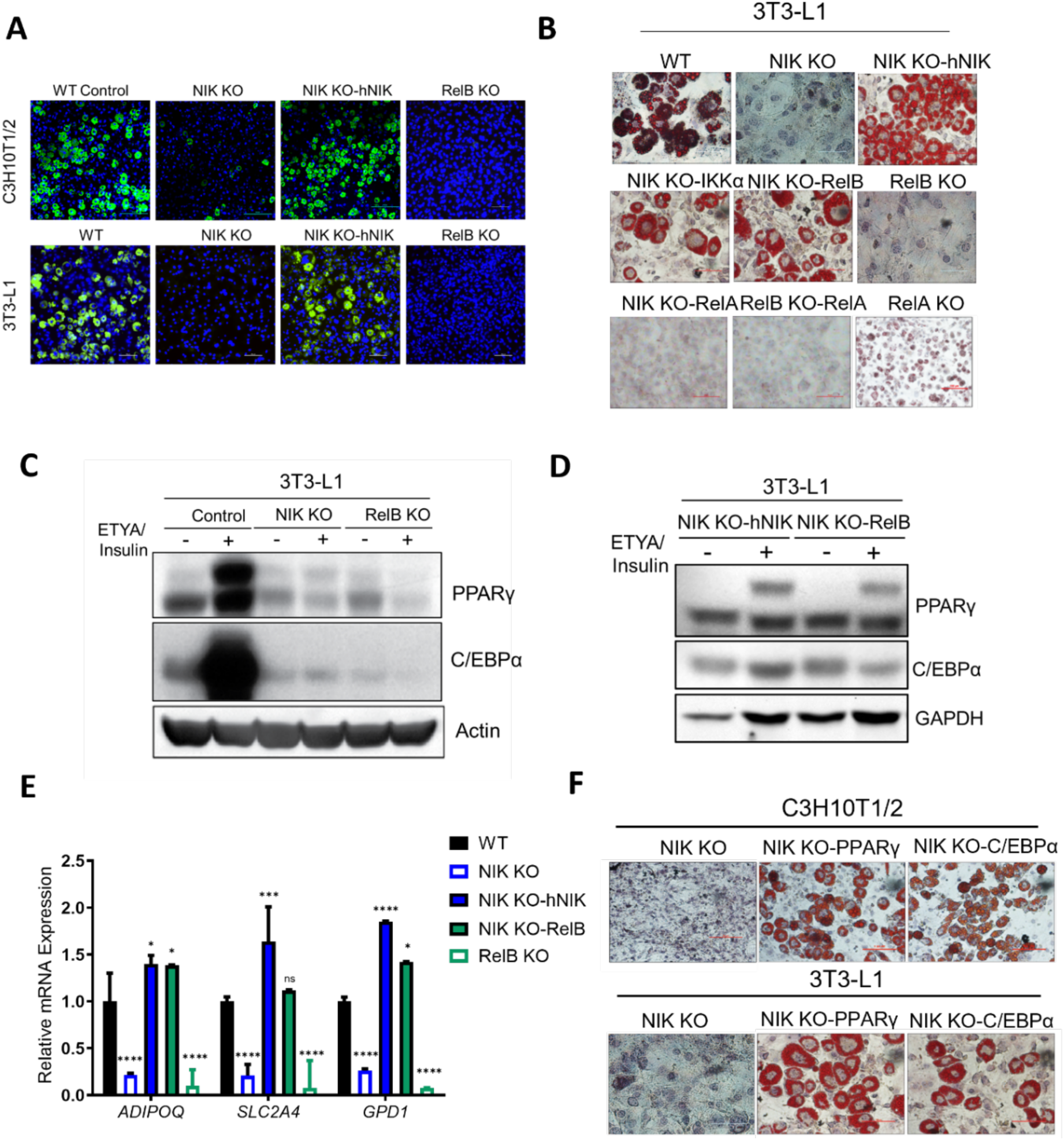
A role for noncanonical NF-κB-dependent signaling in promoting adiposity. **(A)** Fluorescent BODIPY lipid staining of C3H10T1/2 and 3T3-L1 cells showing a significant reduction in adipogenesis with the loss of NIK which is rescued in NIK KO-hNIK cells. **(B)** Oil Red O staining of differentiated 3T3-L1 cells lacking or over expressing various noncanonical/canonical NF-κB proteins. **(C)** PPARγ and C/EBPα expression in undifferentiated vs adipogenic treated in NIK KO and RelB KO 3T3-L1 cells compared to WT control cells. **(D)** PPARγ and C/EBPα expression in undifferentiated vs adipogenic treated 3T3-L1 cells with re-expression of human NIK or overexpression of RelB in NIK KO cells (NIK KO-hNIK, NIK KO-RelB). **(E)** qPCR analysis of adipocyte genes (*ADIPOQ, SLC2A4, GPD1*) in ETYA/insulin treated C3H10T1/2 cells. Data represented as mean ± SD, Tukey’s Multiple Comparison Test. **(F)** Oil Red O staining of differentiated NIK KO or NIK KO with overexpression of PPARγ or C/EBPα in C3H10T1/2 and 3T3-L1 cells.

Next, we evaluated if NIK regulation of adipogenesis required downstream NF-κB signaling. We observed that RelB KO 3T3-L1 cells exhibited impaired adipocyte differentiation and overexpression of IKKα and RelB in NIK KO cells (NIK KO-IKKα, NIK KO-RelB) restored adipocyte differentiation similar to NIK (NIK KO-hNIK). In contrast, overexpression of the canonical NF-κB transcription factor RelA in NIK KO or RelB KO cells was unable to restore adipocyte differentiation and RelA KO cells did not exhibit impaired adipogenesis (Figure 7B, Supp. Figure 6A). Additionally, RelA phosphorylation at Ser536 (p-RelA-S536), a marker of its transcriptional activation, decreased significantly 4-6 days after induction of adipocyte differentiation. Conversely, expression of the noncanonical NF-κB protein RelB, increased throughout adipocyte differentiation (Supp. Figure 6B). Furthermore, activation of the canonical NF-κB pathway with TNFα inhibited adipocyte differentiation in WT C3H10T1/2 cells, whereas preferential activation of the noncanonical NF-κB pathway with TWEAK did not inhibit adipogenesis (Supp. Figure 6C,D).

Next, we observed that NIK and RelB were required for the induction of the essential adipogenic transcription factors PPARγ and C/EBPα [40] (Figure 7C, Supp. Figure 6E). Ectopic expression of NIK or RelB in NIK KO cells rescued their ability to respond to the PPARγ agonist ETYA (eicosatetraynoic acid), and increased PPARγ and C/EBPα expression (Figure 7D, Supp. Figure 6F). PPARγ was also observed to be decreased in NIK KO liver extracts (Supp. Figure 6G). Additionally, expression of the PPARγ-regulated, adipocyte-specific genes adiponectin (*ADIPOQ*), glucose transporter 4 (GLUT4, *SLC2A4*), and glycerol-3-phosphate dehydrogenase (*GPD1*), were significantly impaired in NIK KO and RelB KO cells, basally and in response to a PPARγ agonist. Expression of PPARγ and adipogenic genes, as well as functional adipocyte differentiation was restored in NIK KO cells by re-expression of NIK or RelB (Figure 7E). Additionally, the overexpression of PPARγ or C/EBPα rescued adipocyte differentiation in NIK KO C3H10T1/2 and 3T3-L1 cells, demonstrating that NIK KO cells fail to undergo adipogenesis due to their inability to increase expression of these key adipogenic transcription factors (Figure 7F). Taken together, these findings indicate that NIK promotes adiposity through noncanonical NF-κB/RelB-dependent regulation of PPARγ and C/EBPα expression.

## 3. Discussion

Maintenance of energy balance is critical for normal physiological processes and overall health status. In this study, we demonstrate that NIK is required to for metabolic homeostasis through control of mitochondrial respiration and fitness, establishing a previously unrecognized role for NIK in controlling cellular, tissue, and systemic metabolism in response to nutritional stress. Consistent with *in vivo* findings, *in vitro* and *ex vivo* analysis revealed that NIK KO preadipocytes and tissue exhibited impaired mitochondrial spare respiratory capacity and elevated proton leak. Furthermore, we observed a compensatory upregulation of glycolysis in the absence of NIK to meet energy demands (see Figs. 2, 3 & 6). This metabolic reprogramming has parallels to the Warburg effect in cancer cells [41], which facilitates rapid generation of ATP using glucose as a carbon source for anabolic pathways while controlling ROS production and redox homeostasis [42]. Our results are also consistent with the use of glucose and glycolysis to reduce ATP input for substrate catabolism, especially in a stressed state [29–31].

A key finding of our study is our observation that NIK-deficient exhibit higher energy expenditure resulting in decreased adiposity, even under the chronic stress of overnutrition caused by a HFD, which has been shown to induce oxidative stress and promote mitochondrial dysfunction. We note that NIK KO mice were housed similarly to other immunodeficient animals and analyzed at 2-4 months of age when they appeared healthy and exhibited no sign of infection or dermatitis (see Figure 5A). Our data are consistent with a previous report demonstrating that NIK activity is increased in the livers of genetic (*ob/ob)* or HFD-fed obese mice, and liver-specific loss of NIK increases glucose metabolism [16, 17]. Similarly, another group demonstrated that NIK overexpression in pancreatic islet cells decreases insulin secretion and glucose homeostasis [43], consistent with the increased glucose and insulin tolerance found in our NIK KO mice. In this study, we analyzed adipose tissue as a key metabolic tissue that impacts local and systemic metabolism. While male and female NIK KO mice exhibited reduced adiposity and elevated metabolic demand, only females exhibited systemic metabolic changes under a chow-fed diet, including increased energy expenditure and temperature. Elevated thermogenesis is consistent with increased mitochondrial proton uncoupling due to higher proton leak observed in NIK KO cells, as well as the increased brown adipose deposition in NIK KO mice (see Fig 4). Notably, when challenged with a HFD, both genders exhibited a significant increase in energy expenditure and higher glucose oxidation than WT mice, demonstrating that with overnutrition, glycolytic adaptation is fueled to support the increase in energetic demand. Dimorphic sex differences in energy expenditure might be attributed to distinct hormone-dependent metabolic or immune changes between males and females. For example, although females are more prone to adipose deposition than males, estrogen increases insulin sensitivity thereby protecting female mice from diet-induced obesity [44–46], and estrogen also enhances mitochondrial function in adipocytes [47]. Furthermore, a recent study demonstrated that RANKL/NIK signaling induced postmenopausal obesity in ovariectomized Nik^aly/aly^ mice, which exhibited reduced adipocyte hypertrophy and reduced lipid accumulation, consistent with our findings that NIK promotes adiposity and adipose development [48].

Interestingly, our findings revealed that NIK regulates adiposity through both NF-κB-independent and NF-κB-dependent mechanisms. Notably, NIK regulates mitochondrial dynamics and metabolism independently of downstream NF-κB RelB (see Figure 2). However, it promotes adipocyte differentiation in noncanonical NF-κB-dependent manner, whereas canonical NF-κB activity suppresses adipogenesis (see Figure 7). These observations are consistent with previous data showing that activation or overexpression of the canonical NF-κB/RelA pathway inhibits adipogenesis, and that RelA itself is inhibited by adipogenic-promoting PPARγ agonists *in vitro* and *in vivo* [49, 50]. Furthermore, one study demonstrated an increase of RelB and p52 during adipocyte differentiation [51], while a separate study demonstrated that mice with adipose-specific knockout of Tank-binding kinase 1 (TBK1), a repressor of NIK stability, exhibited a reduction in adipose tissue and adipocyte size under a prolonged HFD [52]. However, a cooperative role for the canonical and noncanonical NF-κB pathways in promoting adipocyte differentiation was previously reported in a study of adipose-specific RelB knockout mice in response to lymphotoxin-β-receptor (LTβR) activation [53]. Our results suggest that there are likely signal-specific roles for RelB in regulating PPARγ expression and adipogenesis. Moreover, NIK may have RelB-independent, or non-cell autonomous effects on adipogenesis and adipose development. In the context of these findings, the signals responsible for activating NIK during adipocyte development and expansion under a HFD warrant further study.

Further investigation is needed to understand if NIK has adipose cell-intrinsic effects *in vivo*, as well as assess how other tissue types, such as skeletal muscle which is a major glucose sink [54], regulate metabolic homeostasis *in vivo*. For example, NIK may impact systemic metabolism indirectly through its regulation of immune functions, as changes in bacterial flora or viral infection, which activate NIK/NF-κB signaling, can alter host metabolism to support replication [55, 56]. Furthermore, as a regulator of lymphoid development, NIK deficiency might affect the metabolic status and recruitment of adipose tissue immune cells that can play critical roles in regulating tissue homeostasis [14, 57]. We note that *in vitro* preadipocytes exhibited metabolic phenotypes similar to *ex vivo* adipose tissue (see Figs. 2 & 3), demonstrating that NIK has similar adipocyte cell-intrinsic and tissue-specific metabolic functions.

Our analysis of NIK knockout mice provides new insight into the role of metabolic dysfunction in the profoundly debilitating phenotypes of primary immunodeficiency diseases (PIDs). Several genetic defects in the NF-κB signaling pathway are associated with PIDs [58–62], including recently described patients with loss-of-function mutations in NIK [10, 11]. Patients with PIDs have been observed to have growth defects, mainly in children [63, 64]. Risk for infection have also been linked to factors including body mass index and adipose deposition due to the immunomodulatory effects of the tissue, but less is understood of the metabolic dynamics in patients with PIDs [65, 66]. An intimate association between primary immunodeficiencies and primary metabolic diseases is supported by the observation that mitochondrial disease patients manifest significant immunological defects, [67, 68], and some studies have linked inborn errors of metabolism (IEM) as mimicking or exacerbating immune defects [69]. Overall, our findings are consistent with the increasing appreciation that mitochondrial functions are important for integrating metabolic cues and maintaining systemic metabolic health. Moreover, our work highlights an important role for NIK in regulating metabolic homeostatic and adaptive adipose remodeling mechanisms in response to a variety of disease contexts, including chronic stress due to aging, immunodeficiency or overnutrition.

## Abbreviations

NIK-NF: κB-Inducing Kinase
NF: κB-Nuclear factor kappa light chain enhancer of activated B cells
IKK⍺: inhibitor of kappa B kinase alpha
PID: primary immunodeficiency
SRC: spare respiratory capacity
OCR: oxygen consumption rate
ECAR: extracellular acidification rate
ingWAT: inguinal white adipose tissue
DEXA: dual energy X-ray absorptiometry
gWAT: gonadal white adipose tissue
RER: respiratory exchange rate
HFD: High-fat diet
PPARγ: peroxisome proliferator activated receptor
C/EBP⍺: CCAAT enhancer binding protein alpha
ETYA: eicosatetraynoic acid
TNF⍺: tumor necrosis factor alpha
TWEAK: tumor necrosis factor weak like.

## Author Contributions

Conceptualization: K.M.P., D.W.L., R.S.; Data Acquisition: K.M.P., D.W.L., J.K.; Writing—original draft preparation, K.M.P.; Writing—review and editing, K.M.P. and R.S.; Visualization: K.M.P., R.S.; Funding acquisition: R.S.; Supervision: R.S. All authors have read and agreed to the publication of the manuscript.

## Acknowledgments

We thank Dr. Joseph M. Rutkowski for 3T3-L1 cells, Dr. Larry Suva and Dr. Kirby for assistance with DEXA imaging, Dr. David Threadgill and his lab, including Dr. Alexandra Trott and Orion Hicks at the Texas A&M Rodent Preclinical Phenotyping Core, for assistance with the EchoMRI^TM^ machine and TSE Phenomaster^TM^ metabolic cages, and Texas A&M Health Science Center COM-CAF core for use of the Seahorse machine.

## 4. Materials and Methods

### 4.1 Animal Procedures & *Ex Vivo* Work

All animal experiments were done in accordance with animal use protocol (2019-0102) with approved IACUC guidelines. *Map3k14* mice were purchased from Jackson lab (B6N.129-Map3k14tm1Rds/J) and maintained by heterozygous breeding. Mice are housed on hypoallergenic, alpha dry bedding with weekly cage changes to minimize dermatitis. Chow diet contained 4% fat while HFD contained 45% fat (Lab Supply 58125). Mice analyzed on a HFD were weaned from adult females on a HFD and then maintained on a HFD for at least 2 months.

### 4.2 Cell Culture

Mesenchymal stem cell line and preadipocyte cell line, C3H10T1/2 and 3T3-L1 were cultured in DMEM supplemented with 10% FBS, 100 U/ml penicillin and 0.1mg/ml streptomycin (Thermo Fisher Scientific, Waltham, MA). 293-T cells were obtained from ATCC (www.atcc.org) and cultured in DMEM supplemented with 10% FBS, 100 U/ml penicillin and 0.1mg/ml streptomycin.

Bone marrow MSC cells were isolated from the femurs of mouse and cultured in DMEM supplemented with 10% FBS, 100 U/ml penicillin and 0.1mg/ml streptomycin.

### 4.3 Bone Marrow Cell Isolation

Mouse femurs are collected and sterilized in 70% ethanol and then were crushed in sterile PBS in a mortar with a pestle. From the crushed femurs, 5mL of DMEM (with 10% FBS and 1% penicillin/streptomycin) is then used to collected cells into a conical tube through a 70μM filter. Cells are then centrifuged at 1500rpm for 8 minutes. Media is removed and the cell pellet is resuspended in 3mL of ACK lysis buffer (Lonza 10-548E) for 2 minutes. ACK cell suspension is diluted with 10mL of media and pelleted again. The cell pellet is resuspended in 10mL of media and filtered through a 40μM filter.

### 4.4 CRISPR-Cas9 Gene Knockout

Oligos encoding guide RNAs for murine NIK and RelB are found in Supplemental Table 1. Each gRNAs were cloned into Lenti-CrispR-v2 (Addgene, Cambridge, MA) respectively. C3H10T1/2 and 3T3-L1 cells were transduced with a mixture of lentiviruses (described below) carrying the three murine gRNAs. Puromycin resistant single clonal cells were isolated by serial dilution and experiments were repeated with at least two clones. Loss of NIK or RelB expression was confirmed by immunoblot analysis. For controls, cells were transduced with empty LentiCrispR-V2.

### 4.5 Lentivirus Production

Lentiviral construct of over-expression constructs (NIK, IKKα, RelB, RelA, PPARγ and C/EBPα) was obtained from DNASU (Tempe, AZ). 24μg of lentiviral plasmids and 72μg of polyethyleneimine (Sigma-Aldrich) were used to transfect 293T cells. After 3 days of transfection, viral supernatant was concentrated 20 fold, to 500μl using Lenti-X Concentrator (Clontech, Mountain View, CA), and 100μl of concentrated virus was used to infect cells. Stably transduced cells were selected for 72hr in medium containing 0.6μg/ml puromycin or 6μg/ml blasticidin (Thermo Fisher).

### 4.6 Seahorse Extracellular Flux Analysis

#### 4.6.1 3T3-L1 Cells

Metabolic activity was analyzed using a Seahorse XFe96 Analyzer (Agilent, Santa Clara, CA). For 3T3-L1 cells, forty-thousand cells per well were plated in the collagen-coated (40ng/μL) Agilent Seahorse XFe96 microplates day before analysis. Mitochondrial Stress Tests were conducted according to the manufacturer’s guidelines. Base media was supplemented with 25mM glucose (Sigma, G7021, St. Louis, MO), 2mM glutamine (Sigma, G85420, St. Louis, MO), and 1mM pyruvate (Gibco). Inhibitors were used at the following final concentrations (10x inhibitor was added to injection ports to reach final concentration): 1μM oligomycin A (Sigma, 75351, St. Louis, MO), 2μM FCCP (Sigma, C2920, St. Louis, MO), and mixture of 0.5μM rotenone (Enzo Life Sciences, Farmingdale, NY, ALX-350-360) and 0.5μM antimycin A (Sigma, A8674, St. Louis, MO). For Glycolysis Stress Tests, cells were glucose-starved for 1hr in Seahorse DMEM base media with 2mM glutamine before analysis. Reagents for injections were used at the following final concentrations (10x reagent was added to injection ports to reach final concentration): 10mM glucose, 10uM oligomycin A, 50mM 2-DG (Thermo Fischer Scientific, 50-519-066). After Seahorse analysis, DNA content was measured using DRAQ5 staining (Thermo Fisher Scientific, 50-712-282) for normalization. Assay was ran in 3 biological replicates with samples ran in 5-8 replicates per assay. Analyses were conducted using Seahorse Wave Controller Software v2.6 and XF Report Generators (Agilent Technologies).

#### 4.6.2 *Ex Vivo* Adipose Tissue

Analysis of metabolic activity of adipose tissue was adapted from [70]. 96 well Seahorse plates were coated twice with Cell-Tak (50μg/mL, Corning® Cell-Tak™ Cell and Tissue Adhesive, Cat. No. 354240). Tissue was excised fresh, rinsed well in PBS, and cut into small, >1mg, pieces of similar size and plated into the center of the wells. For Mitochondria Stress Test, base media was supplemented with 25 mM glucose, 2 mM glutamine, and 1 mM pyruvate. Inhibitors were used at the following final concentrations (10x inhibitor was added to injection ports to reach final concentration): 20μM oligomycin A, 20μM FCCP, and mixture of 20μM rotenone and 20μM antimycin A. Tissue samples are ran in 5-10 replicates per tissue type per assay. Seahorse Analyses were done following Agilent guidelines for the mitochondria stress test [71] and energy phenotyping [72].

For Glycolysis Stress Test base media was supplemented with 2mM glutamine, and reagents for injections were used at the following final concentrations (10x reagent was added to injection ports to reach final concentration): 25mM glucose, 20μM oligomycin A, 100mM 2-DG. Run time for tissue was increased to 30 minutes per injection with four measurements per compound (4 cycles of 3 min mix, 1.5 min wait, 3 min measure). Analyses of glycolysis stress test was done following Agilent guidelines [73]. Afterwards, tissue was homogenized, lysed, and normalized by protein.

### 4.7 Immunofluorescence

For immunofluorescence of C3H10T1/2 mitochondria staining, cells were transfected with 1μg DsRed2-Mito-7 (Mito-dsRed) (Addgene Plasmid #55838) by lipofectamine (Invitrogen L3000008) for 48-72hrs or stained with COXIV 1:200 overnight. Cells were then fixed with 4% PFA for 20 minutes at 37°C. After fixation, cells were washed with PBS and then stained with a fluorescent secondary and/or Hoechst (1:1000) diluted in 1% BSA, .1% Triton-X in PBS. Cells were imaged by confocal (Nikon TI A1R inverted confocal microscope).

### 4.8 BODIPY Staining

For immunofluorescence imaging, BODIPY™ 490/509 (4,4-Difluoro-1,3,5,7,8-Pentamethyl-4-Bora-3a,4a-Diaza-s-Indacene) (Sigma Aldrich 790389) was used. 5mM of BODIPY dye was diluted 1:2500 in PBS and incubate on live cells for 10 minutes at 37°C. Afterwards, cells were washed twice with PBS and then fixed in 4% PFA for 20 minutes at 37°C. After fixation, cells were washed twice with PBS and then stained with Hoechst (1:1000) diluted in 1% BSA, .1% Triton-X in PBS for 10 minutes. Cells were imaged by confocal (Nikon TI A1R inverted confocal microscope).

### 4.9 Lipolysis

About 2-5mg was placed in a 12 well plate in duplicate or triplicate. Tissue was left to sit in 300μL DMEM without FBS for an hour in the incubator. Media was removed for t=0 and isoproterenol was added for a final concentration of 100μM. Tissue was incubated with the isoproterenol for an hour and then media was removed for t=1 timepoint, final samples were taken after overnight incubation. 10μL of each sample at the different points were mixed with 160μL of glycerol reagent (Sigma, F6428) and read at an absorbance of 540nm for glycerol levels.

### 4.10 Adipocyte Differentiation

Adipocyte differentiation was induced by treating confluent cells for 2 days with 5 mg/ml insulin (Thermo Fisher Scientific) and 10μM ETYA (Santa Cruz Biotech, Dallas, TX) in Dulbecco’s modified Eagle’s medium containing 10% FBS, and then every 2 days for 6 Days with insulin (5 mg./ml). Treatment with NIK inhibitor B022 was purchased from MEDCHEM EXPRESS (Cat. No. HY-120501) and used at a concentration of 5μM, TNFα (ProSpec CYT-223) and TWEAK (PeproTech 31006) were used at 10ng/mL.

### 4.11 Oil Red O Staining

For the Oil Red O staining, Oil Red O staining solution (0.5% Oil-Red O in isopropyl alcohol solution-distilled water [60:40]) was filtered through the Whatman no. 1 filter paper. Cells were fixed with 10% formaldehyde solution for 30 min at 37°C, then washed with 60% isopropyl alcohol followed by staining with the filtered Oil Red O solution for 20 min and then washed with distilled water three times. To measure Oil Red O staining of adipocyte differentiated cells, dye is eluted with 100% isopropanol for 10 minutes. Eluted dye is transferred to 96 well plate and optical density measurements were done on Perkin Elmer plate reader at 500nm, .5 sec.

### 4.12 Immunoblot

Cells were lysed in RIPA lysis buffer (Thermo Fisher Scientific) with protease/phosphatase inhibitor cocktail (Thermo Fisher Scientific). Protein was mixed with NuPage 4X LDS sample buffer (Thermo Fisher Scientific) containing .1M DTT and denatured at 100°C for 7 min. Proteins were separated on 8% ∼ 12% SDS-PAGE and transferred to nitrocellulose membranes (Amersham). The membranes were blocked for 1h with 5% non-fat dry milk in 0.1% Tween-20/TBS (TBST) and incubated with primary antibodies diluted in blocking buffer at 4°C overnight. After washing in TBST, membranes were incubated with secondary in BSA for 1 hour at room temperature. The blots were washed with TBST and developed using Chemiluminescent HRP Substrate (EMD Millipore) on ChemiDoc MP Imaging System (Bio-Rad) for detection of HRP or an Odyssey Infrared Imaging system (LI-COR Biosciences) for detection of IRDye fluorescent dyes.

### 4.13 Antibodies

Following antibodies were used C/EBPα (CST8178) (Cell Signaling Technology), COXIV (CST11967s), GPD1 (sc-376219) (Santa Cruz Biotechnology, Dallas, TX), IKKα (CST2682), NFKB2 (CST4882), NIK (CST4994), p-RelA (p-p65) (CST3033), RelA (p65) (sc-8008), PPARγ (CST2443), RelB (CST4992), GAPDH (sc137179), and β-actin (sc69879).

### 4.14 RNA Isolation, cDNA Synthesis, and Quantitative-RT-PCR

Primers used for qPCR are in Supplemental Table 2.

#### 4.14.1 Cells

Total RNA was isolated from cells by Purelink™ RNA Mini Kit (Life Technologies). cDNA was synthesized from 1 μg total RNA using iScript reverse transcription supermix (Bio-Rad, Hercules, CA) following the manufacturer’s protocol. Quantitative RT-PCR was performed using iTaq Universal SYBR Green Supermix (Bio-Rad) with StepOnePlus Real-Time PCR System (Applied Biosystems, Foster City, CA).

#### 4.14.2 Tissue

Mouse tissue was frozen with liquid nitrogen and ground to a powder with a pestle in a mortar (a minimum of 50mg of tissue is needed, for WAT 1g may be needed). Tissue was then lysed with Trizol (1mL per gram of tissue), followed by addition of chloroform to centrifuged supernatant (200μL chloroform: 1mL Trizol). Equal amounts of 70% ethanol was added to equal parts of aqueous layer from sample in a new tube and then RNA was then collected in subsequent steps using Invitrogen Purelink RNA mini kit. Zymo DNAse was used with 40-80μL per sample. cDNA was synthesized from 2μg of RNA using iScript reverse transcriptase and buffer mix. Samples were ran in triplicate.

### 4.15 EchoMRI^TM^

EchoMRI^TM^ 100H machine was used to analyze fat and lean mass of chow and HFD mice at 2 or 4 months of age. Mice were weighed before imaging and imaged in MRI tubes.

### 4.16 DEXA Imaging

Dual-energy x-ray absorptiometry was used on anesthetize mice to visualize low and high-density tissue. Mice from maintained on either a chow or HFD were used.

### 4.17 Metabolic Cages

Metabolic analysis on the mice was conducted in TSE Phenomaster^TM^ metabolic cages through Rodent Preclinical Phenotyping Core at TAMU (https://genomics.tamu.edu/preclinical-phenotyping/). Mice were weighed the day of and housed individually for 48hrs. Data was analyzed based on last 24hrs to compensate for mouse adjustment to the new environment.

### 4.18 Glucose Tolerance Testing

For glucose tolerance testing mice were morning fasted for 6hrs in clean cages. Mice were weighed, and from tail clip, initial glucose levels are recorded using a glucose meter. The mice are then intraperitoneally injected with 20% D-glucose at 2g/kg (μL= 10 x BW). Blood glucose (in mg/dL) is then recorded at t=15, t=30, t=60, t=120 minutes post injection.

### 4.19 Insulin Tolerance Testing

For insulin tolerance testing mice were morning fasted for 4hrs in clean cages. Mice were weighed and from tail clip initial glucose levels are recorded using a glucose meter. The mice are then intraperitoneally injected with .5 U/kg Insulin (μL= 5 x BW of .1 U/mL insulin). Blood glucose (in mg/dL) is then recorded at t=15, t=30, t=45, t=60, t=90 minutes post injection.

### 4.20 Statistical Analysis

Statistical Analysis was done using GraphPad PRISM software, specifics on data representation and tests used for analysis can be found in figure legends. *p ≤.05, ** p ≤.01, *** p ≤.001, **** p≤.0001. Unpaired student t-tests were ran as two-tailed. All statistically significant analyses were ran based on a 95% confidence interval.

