## Supplementary Data for "NF-κB-Inducing Kinase Maintains Mitochondrial Efficiency and Systemic Metabolic Homeostasis"

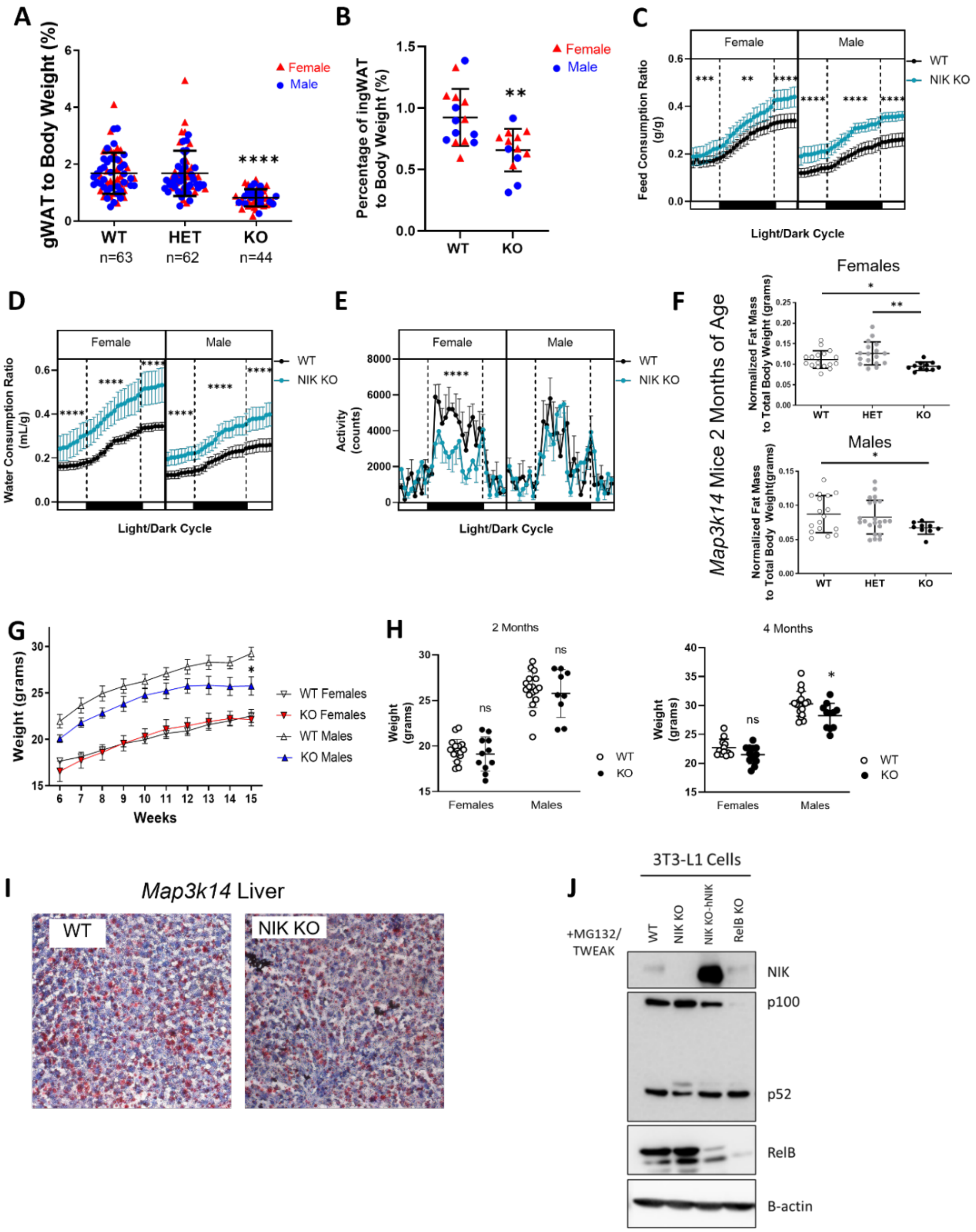

**Supplemental Figure 1.**

**(A)** Weights of 4-month-old chow fed WT, HET, and NIK KO male and female gonadal WAT on a chow diet. Data are represented as mean ± SD, **** p ≤.0001, Tukey Multiple Comparison Test. **(B)** Analysis of inguinal WAT from chow fed male WT and NIK KO mice. Data represented as mean ± SD, ** p ≤.01, Unpaired Student t-test. **(C-E)** Metabolic cage data of male and female WT, HET, NIK KO mice at 4 months of age for **(C)** food consumption **(D)**, water consumption, and **(E)** activity**.** Data are represented as mean ± SEM, *p≤.05, ** p ≤.01, **** p ≤.0001, Unpaired Student t-test. **(F)** Echo MRI data of fat and lean mass ratios of male and female Map3k14 mice at 2 months of age. Data represented as mean ± SEM, * p≤.05, ** p ≤.01, *** p ≤.001, Unpaired Student t-test. **(G)** Average weights of WT and NIK KO mice between males and females from 6-15 weeks (Female *Map3k14* mice; WT n= 9 and KO n=7, Male *Map3k14* mice; WT n=6 and KO n=5). Data represented as mean ± SEM, Sidak's Multiple Comparisons Test. **(H)** Weights from chow-fed, male and female WT and NIK KO mice at 2 months and 4 months of age. Data are represented as mean ± SD, 2-month-old Males; WT= 17, KO=9, Females; WT= 17, KO=11, 4-month-old Males; WT=18, KO=11, Females; WT= 12, KO=12, Unpaired Student t-test. **(I)** Oil Red O and hematoxylin staining of liver sections from chow fed WT and NIK KO mice. **(J)** Verification immunoblot of MG132 (10μM) and TWEAK(10ng/mL) treated 3T3-L1 cells with CRISPR-Cas9 modifications for knockout of NIK or RelB, and rescue of human NIK in the NIK KO cell line.

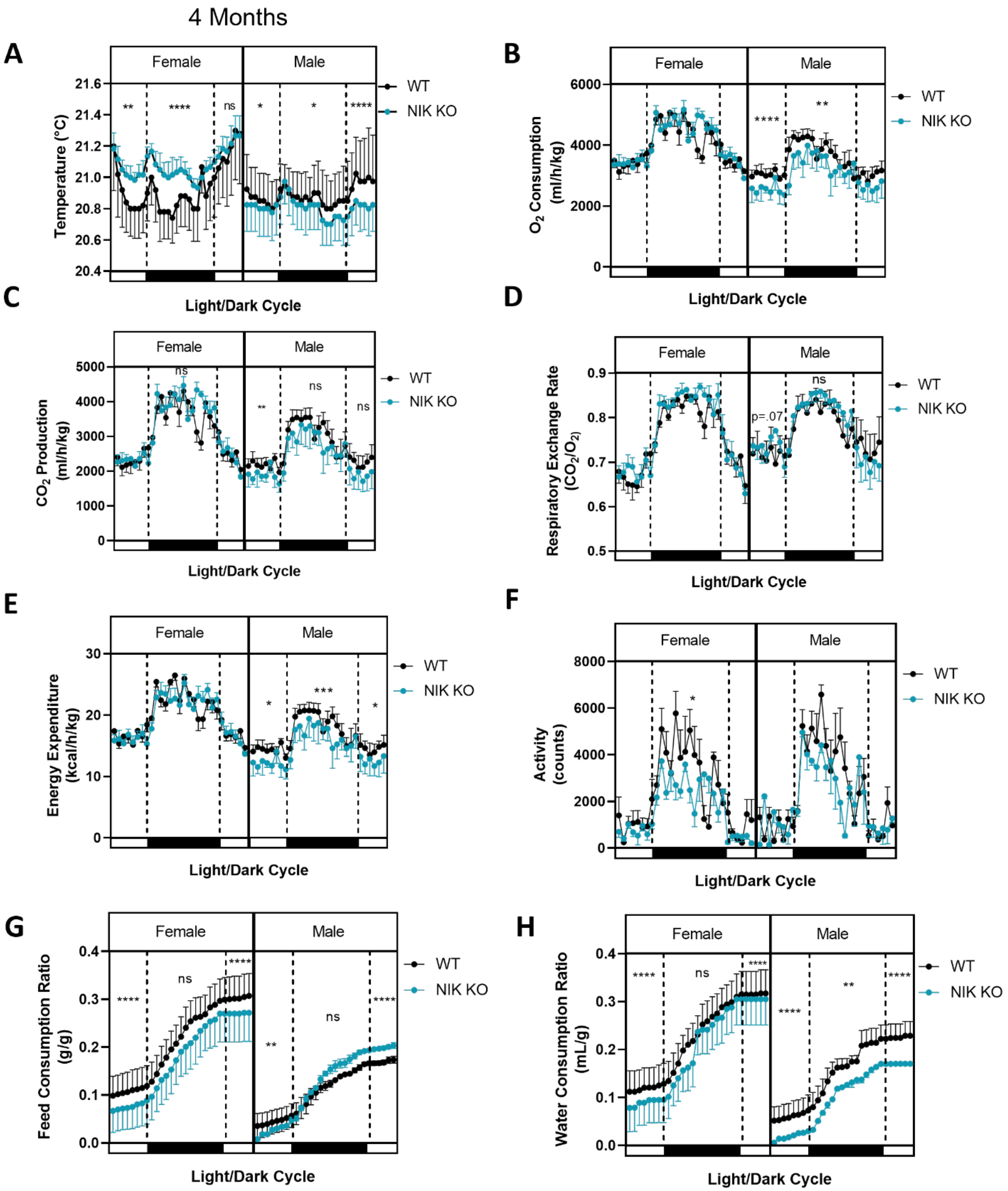
4 Months
A
Female
Male
WT
NIK KO
Temperature (°C)
21.6
21.4
21.2
21.0
20.8
20.6
20.4
**
****
ns
*
*
****
Light/Dark Cycle
B
Female
Male
WT
NIK KO
O2 Consumption (ml/h/kg)
6000
4000
2000
0
****
**
Light/Dark Cycle
C
Female
Male
WT
NIK KO
CO2 Production (ml/h/kg)
5000
4000
3000
2000
1000
0
ns
**
ns
ns
Light/Dark Cycle
D
Female
Male
WT
NIK KO
Respiratory Exchange Rate (CO2/O2)
0.9
0.8
0.7
0.6
0.5
p=.07
ns
Light/Dark Cycle
E
Female
Male
WT
NIK KO
Energy Expenditure (kcal/h/kg)
30
20
10
0
*
***
*
Light/Dark Cycle
F
Female
Male
WT
NIK KO
Activity (counts)
8000
6000
4000
2000
0
*
Light/Dark Cycle
G
Female
Male
WT
NIK KO
Feed Consumption Ratio (g/g)
0.4
0.3
0.2
0.1
0.0
****
ns
****
**
ns
****
Light/Dark Cycle
H
Female
Male
WT
NIK KO
Water Consumption Ratio (mL/g)
0.4
0.3
0.2
0.1
0.0
****
ns
****
****
**
****
Light/Dark Cycle

**Supplemental Figure 2.**

**(A-H)** Analysis of a 24hr period of metabolic cage data representing chow fed male and female, WT and NIK KO mice **(A)** Metabolic cage temperature of chow fed female and male WT and NIK KO mice at 4 months of age. **(B-H)** Metabolic cage data from 2-month-old *Map3k14* mice. Data are represented as mean ± SEM, Males; WT= 5 KO=3, Females; WT=6, KO= 5, *p≤.05, ** p ≤.01,***p ≤.001, **** p ≤.0001, Unpaired Student t-test.

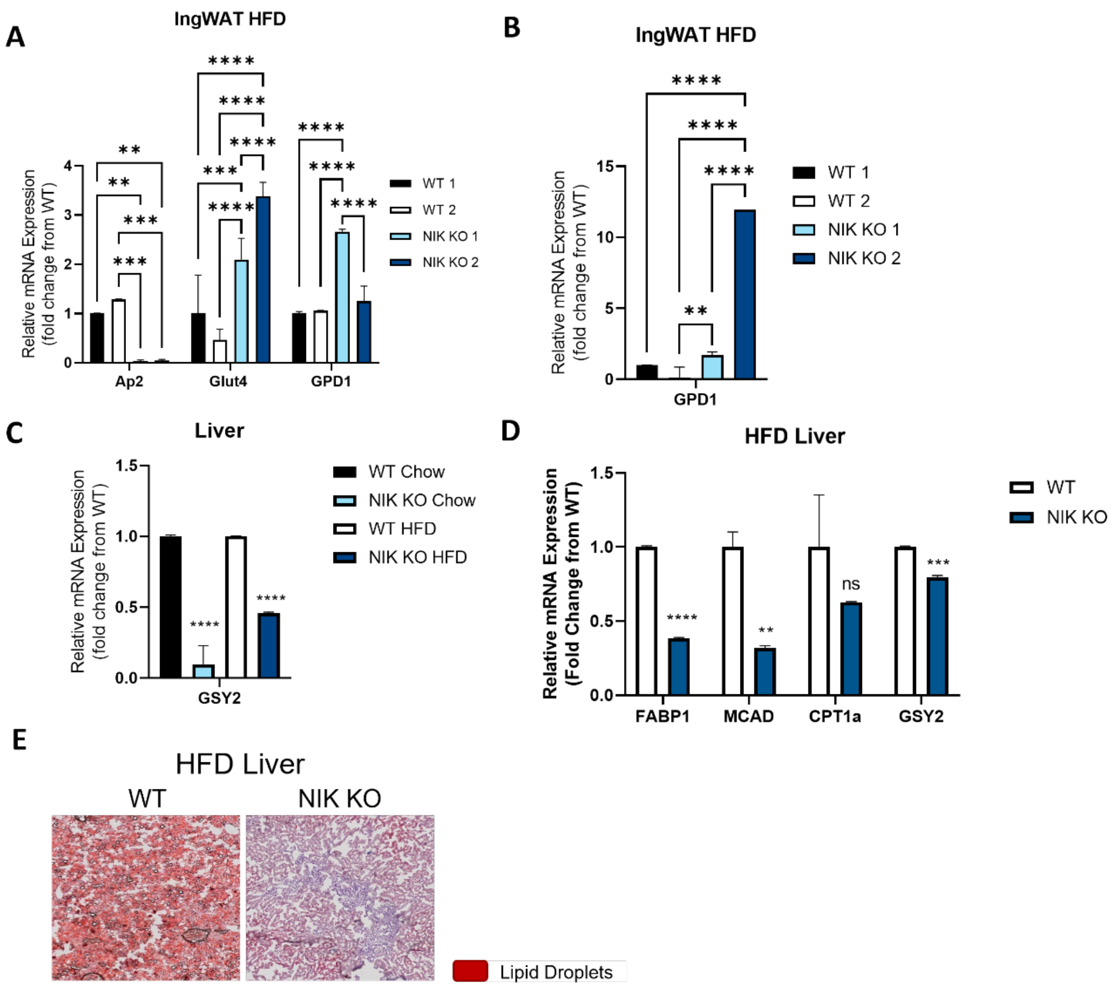

**Supplemental Figure 3.**

**(A)** qPCR analysis of adipocyte genes Ap2 (adipocyte specific protein 2; FABP4), GLUT4 (glucose transporter 4, SLC2A4), GPD1 (glucose-3-phosphate dehydrogenase 1) of inguinal WAT from male HFD mice. **(B)** GPD1 mRNA analysis of inguinal WAT from HFD female mice. **(C)** GSY2 (glycogen synthase 2) relative mRNA expression from chow and HFD liver samples. **(D)** qPCR analysis of fatty acid oxidation genes (FAB1; fatty acid binding protein, MCAD; medium chain acyl-CoA dehydrogenase, CPT1α; carnitine protein transferase) and glycogen synthase 2 from HFD female liver samples. N=2 biological replicates ran in triplicate **(A-D)** Data represented as mean ± SD, Unpaired Student t-test. **(E)** Oil Red O and hematoxylin staining for lipid accumulation in liver samples of HFD mice.

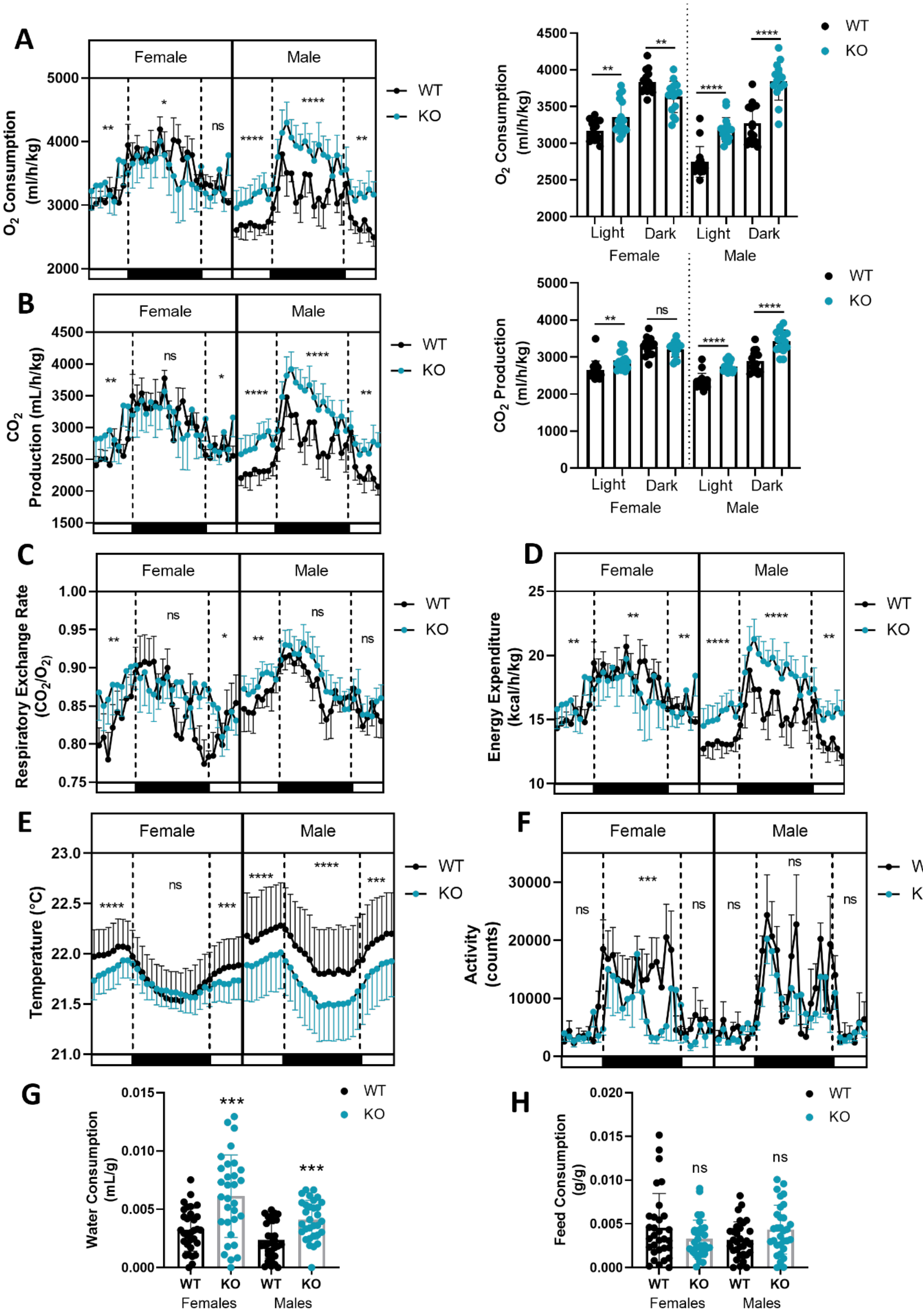
A
B
C
D
E
F
G
H
Female
Male
WT
KO
O2 Consumption (ml/h/kg)
Light
Dark
CO2 Production (mL/h/kg)
CO2 Production (ml/h/kg)
Respiratory Exchange Rate (CO2/O2)
Energy Expenditure (kcal/h/kg)
Temperature (°C)
Activity (counts)
Water Consumption (mL/g)
Feed Consumption (g/g)
Females
Males
ns

**Supplemental Figure 4.**

**(A-H)** Data collected from HFD male and female *Map3k14* mice individually housed in metabolic cages at about 3 months of age. Morning and night analysis from a 24-hour time period of **(A)** oxygen consumption, **(B)** $CO_2$ production, **(C)** respiratory exchange rate ($CO_2$/ $O_2$), and **(D)** caloric energy expenditure, **(E)** metabolic cage temperature, **(F)** activity, **(G)** water consumption normalized to body weight, and **(H)** food consumption normalized to body weight. Line graphs represented as mean ± SEM, bar graphs represented as mean ± SD. Males; WT= 5 KO=8, Females; WT=7 KO=6. Data analyzed by Unpaired Student t-test.

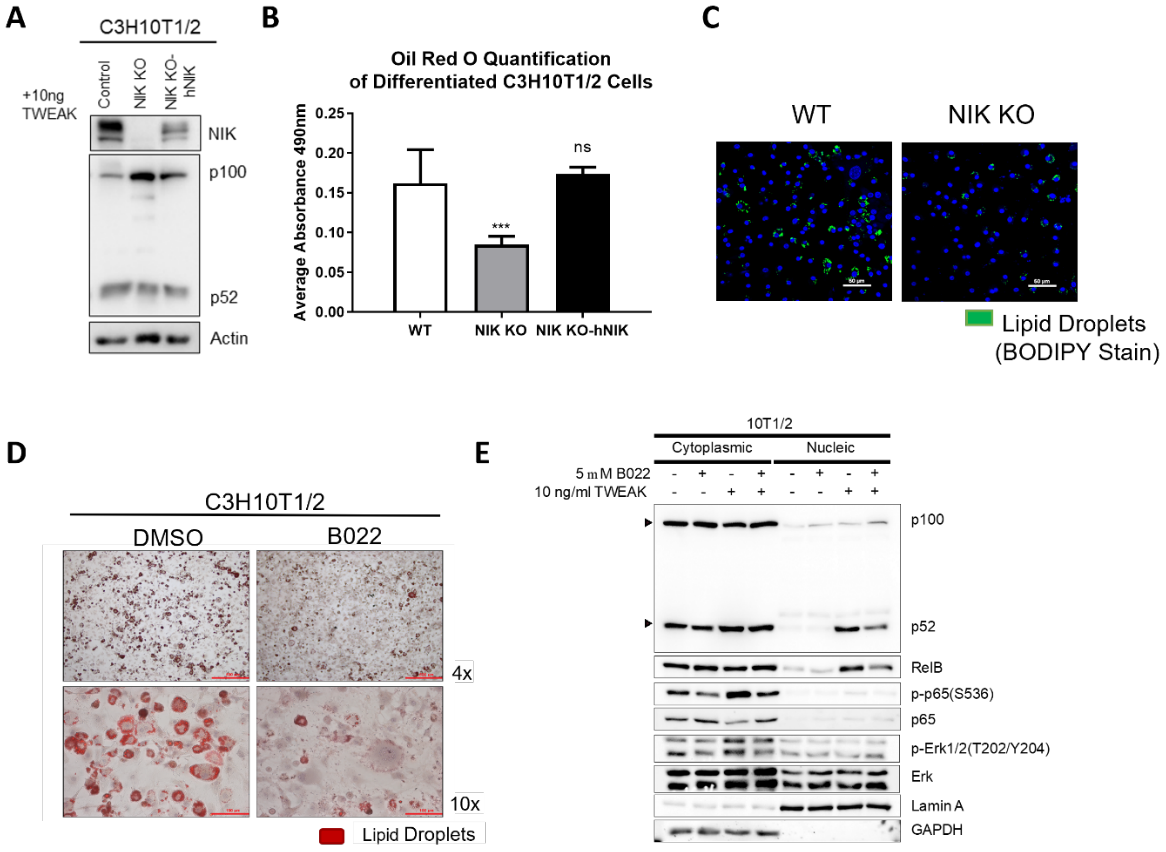

**Supplemental Figure 5.**

**(A)** Blot of C3H10T1/2 cells confirming the loss of NIK (NIK KO) and the rescue of non-canonical NF-κB functionality (p100-p52 processing) with the reestablishment of human NIK (NIK KO-hNIK). . **(B)** Quantification of oil red o staining of differentiated C3H10T1/2 cells. Data represented as mean ± SD, * p≤.05, ** p ≤.01, *** p ≤.001, **** p ≤.0001, Tukey's Multiple Comparison Test. **(C)** Adipocyte differentiation of primary bone marrow cells from WT and NIK KO mice stained with BODIPY (lipid droplets; green) and Hoechst (blue). **(D)** C3H10T1/2 cells treated with DMSO or the NIK inhibitor (B022), induced for adipocyte differentiation, and then stained with Oil Red O. **(E)** Cytoplasmic/ nuclear fractionation of C3H10T1/2 cells treated with B022 (NIK inhibitor) or TWEAK (activator of the noncanonical NF-κB pathway), showing inhibition of p100-p52 processing (induction of noncanonical protein processing by NIK) with B022 treatment.

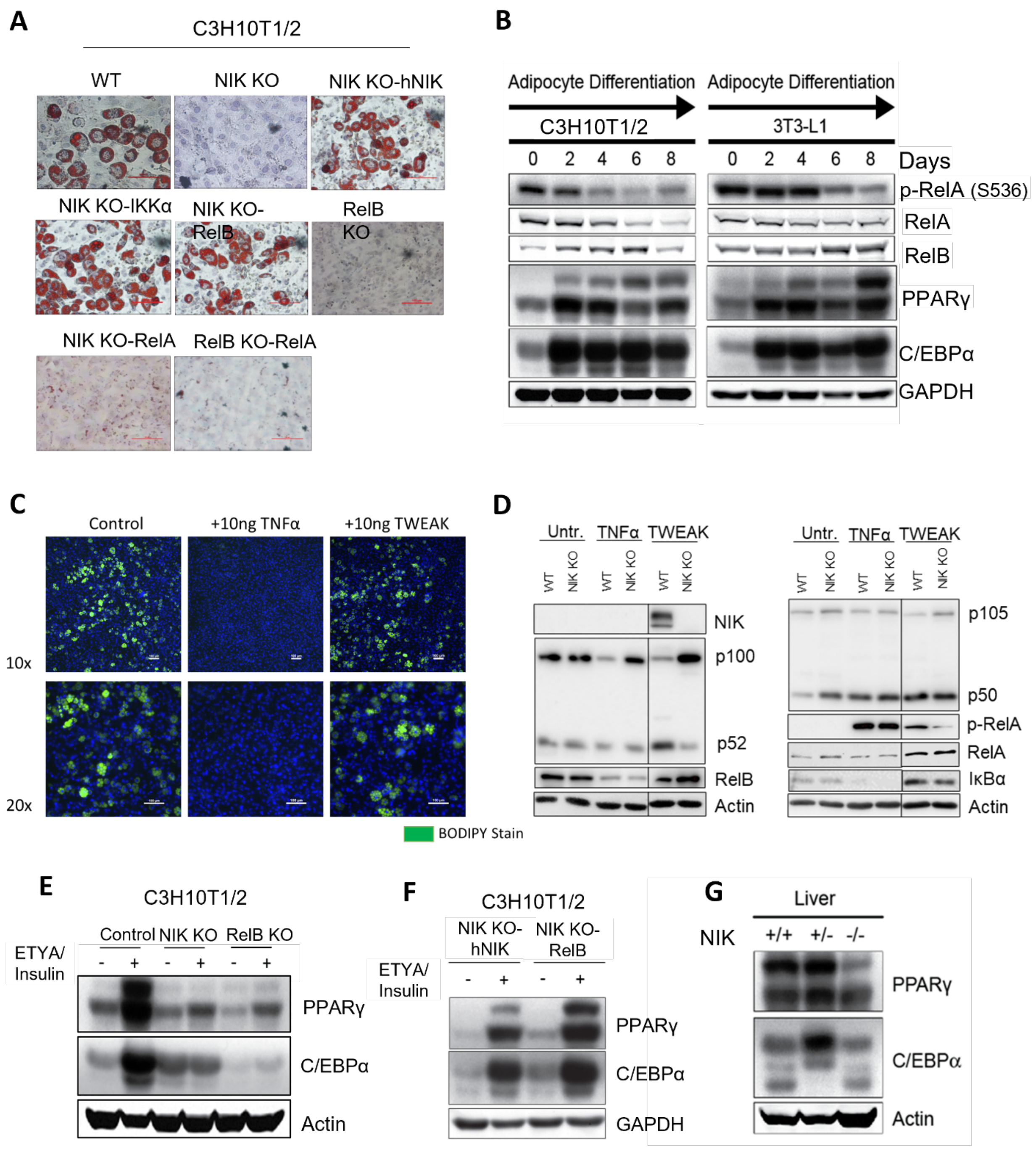

**Supplemental Figure 6.**

**(A)** Oil Red O staining of C3H10T1/2 cells after induction of adipocyte differentiation with ETYA and insulin. **(B)** Immunoblot of adipocyte induction in C3H10T1/2 and 3T3-L1 cells with ETYA/Insulin. **(C)** BODIPY staining of lipid droplets (green) and Hoechst staining (blue) in differentiated C3H10T1/2 treated with TNFα (tumor necrosis factor alpha; activator of canonical NF-κB pathway) or TWEAK (tumor necrosis weak life factor; activator of noncanonical NF-κB pathway). **(D)** Immunoblots confirming activation of the noncanonical or canonical NF-κB pathway after induction with TWEAK or TNFα, respectively. Immunoblot of transcriptional activators of adipogenesis, PPARγ and C/EBPα expression in undifferentiated and differentiated **(E)** NIK KO, RelB KO, **(F)** NIK KO-hNIK and NIK KO-RelB C3H10T1/2 cells, **(G)** and mouse liver samples.

| Target Gene | Guide RNA Sequence |
| --- | --- |
| NIK | gNIK-1: AGGAUGGGGCAUUCCGCUGU |
| | gNIK-2: UACCUGGUGCAUGCGCUCCA |
| | gNIK-3: AGUAUCGAGAAGAGGUCCAC |
| RelB | g RelB-1: AGCGGCCCUCGCACUCGUAG |
| | gRelB-2: GCGCUUCCGCUACGAGUGCG |
| | gRelB-3: ACUGCACGGACGGCGUCUGC |

**Supplemental Table 1:**

Guide RNA sequences used for CRISPR-Cas9 knockout of NIK or RelB in cell culture.

| Target Gene | Forward Sequence | Reverse Sequence |
|---|---|---|
| *AIPOQ* (Adiponectin) | 5'- GTT GCA AGC TCT CCT GTT CC -3' | 5'- GAG CGA TAC ACA TAA GCG GC -3' |
| *AP2* | 5'- GTG CTG CAG CCT TTC TCA C-3' | 5'- GTT CCC ACA AAG GCA TCA C-3' |
| *CPT1α* | 5'-CTG ATG ACG GCT ATG GTG TTT-3' | 5'-GTG AGG CCA AAC AAG GTG ATA-3' |
| *FABP1* | 5'- CCA ATT GCA GAG CCA GGA GA-3' | 5'- CCC CTT GAT GTC CTT CCC TTT-3' |
| *GPD1* | 5'- ACA CCC AAC TTT CGC ATC AC -3' | 5'- TAG CAG GTC GTG ATG AGG TC -3' |
| *GSY2* | 5'- CAC ATC ACC ACC AAC GAC GGA-3' | 5'- TTT AGC CGA TCC CTC TCA GCC-3' |
| *LEP* (Leptin) | 5'- TTC CTG GTG GCT TTG GTC CTA -3' | 5'- AGC ACA TTT TGG GAA GGC AG -3' |
| *MCAD* | 5'-ACC CTG TGG AGA AGC TGA TG-3' | 5'-AGC AAC AGT GCT TGG AGC TT-3' |
| *SLC2a4* (GLUT4) | 5'- GAA ACC CAT GCC GAC AAT GA -3' | 5'- CTG TGC CAT CTT GAT GAC CG -3' |
| *GAPDH* | 5'- CTT TGT CAA GCT CAT TTC CTG G -3' | 5'- TCT TGC TCA GTG TCC TTG C -3' |
| *RPLP0* | 5'-AAG CGC GTC CTG GCA TTG TCT-3' | 5'-CCG CAG GGG CAG CAG TGG T- 3' |

**Supplemental Table 2:**

Primers used for qPCR analysis of cells or tissue. 3T3-L1 gene expression was normalized to GAPDH. Mouse tissue gene expression was normalized to RPLP0.
